# Orthogonal resistance mechanisms of classical- and induced-proximity inhibitors

**DOI:** 10.1101/2025.05.10.652755

**Authors:** Manuel L. Merz, Karishma Kailass, Rajaiah Pergu, Kien Tran, Kritika Gupta, Zachary C. Severance, Sameek Singh, Sreekanth Vedagopuram, Benjamin K. Law, Gideon Rosenblatt, Rohil Dhaliwal, Amit Choudhary

**Author notes:** These authors contributed equally to this work and are listed arbitrarily (Manuel L. Merz, Karishma Kailass). They will put their name first in their curriculum vitae citations or elsewhere. These authors contributed equally to this work and are listed arbitrarily (Rajaiah Pergu, Kien Tran, Kritika Gupta). They will put their name first in their curriculum vitae citations or elsewhere. To whom correspondence should be addressed: Amit Choudhary, Chemical Biology and Therapeutics Science, Broad Institute of MIT and Harvard, 415 Main Street, Rm 3012, Cambridge, MA 02142.

## Abstract

Resistance development is an inevitable failure mode of many drugs, pointing to the need to develop agents with orthogonal resistance mechanisms. Induced-proximity modalities, an emergent class of therapeutics, operate by forming a ternary complex with the protein-of-interest (POI) and effectors, unlike classical inhibitors that form binary complexes with the POI. Using KRAS as a model system, we employed base editor tiling mutagenesis screening to show that induced-proximity inhibitors exhibit orthogonal resistance mechanisms to classical inhibitors despite overlapping binding sites, offering an opportunity to circumvent resistance mechanisms of classical inhibitors. These findings highlight the use of base editor mutagenesis screens to prioritize inhibitors with orthogonal resistance mechanisms and the potential of induced-proximity inhibitors to overcome the drug resistance of classical inhibitors.

---

A contemporary approach to combating drug resistance is to target a different binding pocket on the same protein.^1^ For example, allosteric inhibitors of kinases have been developed for emergent resistance to active-site inhibitors.^2^ However, allosteric sites are often not readily available and challenging to discover, limiting the generalizability of this strategy. An alternative approach could be to target the same binding site but leverage a distinct inhibitory mechanism to overcome drug resistance. Recently, there has been a surge in interest for induced-proximity modalities (e.g., molecular glues) that operate by promoting proximity between the target and an effector protein to form a ternary complex,^3^ which is in contrast with classical inhibitors (e.g., active-site inhibitors) that operate by forming a binary complex with the target protein (**Figure 1a**). We hypothesized that molecular glues and classical inhibitors will have orthogonal resistance mechanisms, even with the same binding site on the target protein, owing to different mode-of-action. We validate this hypothesis using oncogenic KRAS (i.e., KRAS^G12C^ or KRAS^G12D^) as a model and tiling mutagenesis with base editors, a CRISPR-based technology that allows targeted modifications of DNA base at a genomic locus (e.g., C→T, A→G).^4, 5^

**Figure 1.**
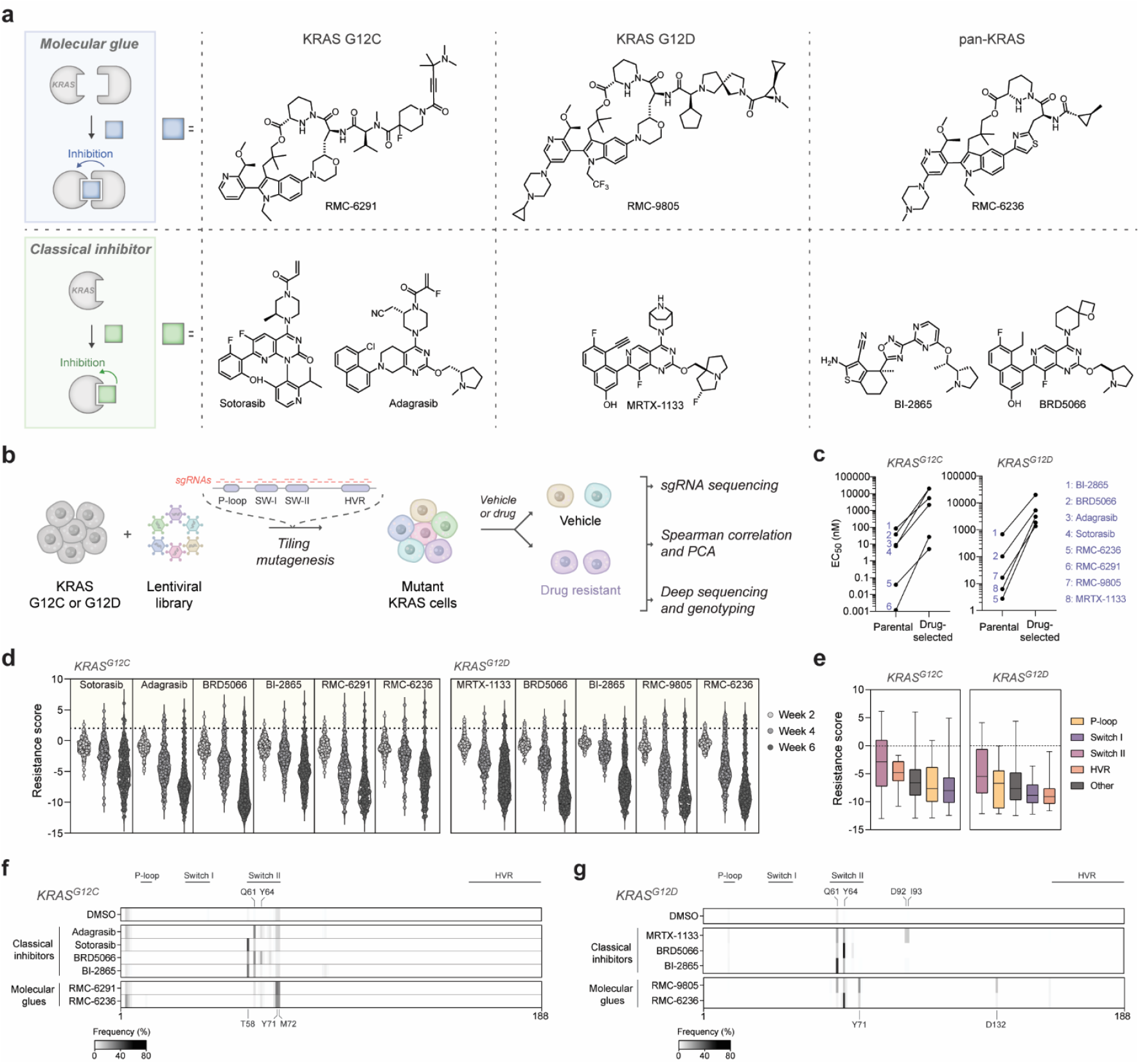
Base editor tiling mutagenesis identifies drug-resistant KRAS^G12C^ and KRAS^G12D^ cells. **a** Chemical structures of KRAS inhibitors. KRAS inhibitors are separated by their modality, forming either a ternary complex as a molecular glue (blue square, top) or a binary complex as a classical inhibitor (green square, bottom). **b** Overview of workflow for tiling mutagenesis using base editors. **c** Pair-wise comparison showing inhibitor sensitivity of different selections in KRAS^G12C^ (left) and KRAS^G12D^ (right) cells transduced with adenine base editor (ABE). Each datapoint shows the mean of EC_50_ across three independent selection replicates. **d** Mutagenesis screen in KRAS^G12C^ and KRAS^G12D^ cells transduced with ABE. The drug dose was escalated from EC_30_ to EC_50_ to EC_80–90_ over 6 weeks (2 weeks per dose) and sgRNA enrichment measured. The dotted line marks log2 fold-change sgRNA enrichment (LFC) = 2, corresponding to >2 SD above the mean of non-targeting controls (n = 78). Each datapoint represents the LFC of a sgRNA across three independent selection replicates. **e** Box plot showing LFC sgRNA enrichment in different KRAS domains in KRAS^G12C^ and KRAS^G12D^ cells transduced with ABE. **f–g** Deep sequencing of alleles corresponding to the coding sequence of KRAS. Each of the four exons on KRAS was PCR-amplified and sequenced individually for all selections performed in KRAS^G12C^ (**f**) and KRAS^G12D^ (**g**) cells. Mutational frequency was calculated as the sum of allele frequencies carrying a missense mutation at the indicated residues.

We chose the oncogenic KRAS and base editor mutagenesis strategy for multiple reasons. First, classical and molecular glue inhibitors of oncogenic KRAS that target the same region (i.e., around switch-II pocket) are readily available with exhaustive mechanistic, structural, and clinical data (**Figure 1a** and **Figure S1a, S1b**).^6^ Second, while classical- and molecular glue-targeting KRAS^G12C^ inhibitors are covalent, those targeting KRAS^G12D^ are non-covalent, affording examination of covalency as another variable. Third, oncogenic KRAS rapidly mutates to develop resistance and is a high therapeutic value target.^7, 8^ Several KRAS inhibitors are in clinical trials, and there is an urgent need to understand possible modes of resistance and alternative treatment options. Fourth, tiling mutagenesis using base editors allows chronic drug treatment for resistance development (> 6 weeks vs. < 2 weeks for saturation mutagenesis studies) that mimics the therapeutic regimen. Fifth, the base editors mutate at the endogenous loci in the native context and expression of oncogenic KRAS.^9^ In contrast, deep mutational scanning platforms overexpress mutant KRAS beyond endogenous levels^10, 11^ that affects protein-protein interactions and causes cellular senescence.^12^ Maintaining endogenous levels of target and effector proteins is particularly important for evaluating proximity-based modalities, as enhancing effective molarities is their primary mode of action.^13^ Finally, base editor tiling mutagenesis primarily introduces transition mutations, which are more likely to emerge as acquired resistance mutations.

We initiated our investigations by assessing the effects on the cell viability of various inhibitors on cell lines harboring mutations at glycine 12, a frequently mutated site in oncogenic KRAS (**Figure S1c**). We selected three molecular glue inhibitors (**Figure 1a**) that stick cyclophilin (CypA) onto KRAS (RMC-6291 for KRAS^G12C^, RMC-9805 for KRAS^G12D^ and RMC-6236 for pan-KRAS).^14-16^ We chose five classical inhibitors (sotorasib and adagrasib for KRAS^G12C^, MRTX-1133 for KRAS^G12D^, BRD5066 and BI-2865 for pan-KRAS).^17-20^ While most inhibitors potently killed cells harboring on-target mutations (e.g., inhibitors for KRAS^G12C^ or KRAS^G12D^ variants potently killed cells harboring respective variants), the potency was substantially reduced (EC_50_ > 1 μM) in cells with non-target KRAS^G12X^ mutations, highlighting that most drugs are one point mutation away from developing resistance (**Figure S1c**). Indeed, the target engagement of these mutant-specific inhibitors relies heavily on covalent bonding with cysteine in KRAS^G12C^ or salt bridge formation with aspartate in KRAS^G12D^.^17^ We envisioned additional hotspots for escape mutations beyond KRAS^G12X^ could confer similar resistance and this motivated us to perform tiling mutagenesis studies.

Cytosine base editors (CBE) allow facile conversion of C to T while adenine base editors (ABE) convert A to G, and these editors can perform tiling mutagenesis (**Figure 1b**) across the gene using barcoded single guide RNA (sgRNA).^9, 21-25^ Using the CRISPOR tool^26^ and the NGN protospacer adjacent motif (PAM) sequence, we designed a sgRNA library across the four exons of KRAS and excluded sequences with predicted high off-target scores against the human genome assembly GRCh38.^27^ Using ABE and CBE on either strand of the DNA, the designed sgRNA library can mutagenize nearly all residues in KRAS (**Figure S2a, S2b**). We cloned the sgRNA library into single plasmid systems, with either CBE (BE3.9max) or ABE (ABE8e),^28^ and delivered them into MIA PaCa-2 (KRAS^G12C^) and AGS (KRAS^G12D^) cells to create a pool of cells carrying diverse KRAS variants. These pools were subjected to drug selection over six weeks by gradually increasing the drug dose from EC_30_ to EC_90_ via EC_50_, eventually enriching for cells harboring escape mutations (and corresponding barcoded sgRNAs that yield such mutations). KRAS^G12C^ cells were treated with sotorasib, adagrasib, and RMC-6291, while KRAS^G12D^ cells were exposed to MRTX-1133 and RMC-9805. Both cell lines were treated with the pan-KRAS inhibitors BRD5066, BI-2865, and RMC-6236, giving a total of eleven drug and one vehicle selection for each base editor.

Several lines of evidence validate our mutagenesis-screening pipeline. First, while the EC_50_ of all inhibitors in the parental cell line were in low nanomolar, a 20–850-fold increase in the EC_50_ was observed in the resistant population for all inhibitors and ABE-transduced KRAS variants (**Figure 1c**). Second, we observed a gradual depletion of cells harboring most sgRNAs with increasing inhibitor concentrations (**Figure 1d**), but an enrichment of sgRNAs targeting the switch II domain, where all the tested inhibitors are known to bind (**Figure 1e** and **Figure S3**). Indeed, deep sequencing of the KRAS coding regions showed a strong accumulation of mutations around switch II pocket residues 58-72 that are directly or indirectly involved in drug-binding through high-affinity side-chain interactions, which aligns with previous observations for base editor screens performed with adagrasib and sotorasib in NCI-H23 cells (KRAS^G12C^) (**Figure S1a, S1b**).^25^ Finally, our screen showed enrichment of clinically relevant Q61 secondary mutations in both KRAS^G12C^ and KRAS^G12D^, which were previously characterized as resistance mutations for adagrasib in patient-derived non-small-cell lung cancers (**Figure 1f, 1g** and **Figure S4c–f**).^7, 29^ We also confirmed mutation sites previously identified in deep mutational scanning, including Y64 and M72 mutations (**Figure 1f, 1g**).^10, 29^

We next investigated the similarities and differences in resistance patterns of molecular glues and classical inhibitors and between KRAS variants. We compared the sgRNA enrichment pattern of molecular glue *versus* classical inhibitors following EC_80_ selection using a principal component analysis (PCA) of ABE transduced cells. Here, we examined the angles between the loadings in the biplot, where smaller acute angles and larger obtuse angles indicate stronger positive and negative correlations, respectively, between variables used in the PCA. We observed very weak correlation between distinct inhibitor classes, as the angles between loadings for KRAS^G12C^ and KRAS^G12D^ variants measured to be 81 ± 13° and 69 ± 2°, respectively (**Figure 2a**). In contrast, inhibitors within a class shared strong correlations for both KRAS^G12C^ and KRAS^G12D^, with measured angles between loadings of 16 ± 12° and 2 ± 1°, respectively. This near-orthogonality is further supported by Spearman correlation analysis of the sgRNA enrichments for the various inhibitors, in which we generally found a poor correlation between classical- and molecular-glue inhibitors, but a positive correlation within inhibitors of the same class (**Figure S5**). To better understand sgRNA enrichment for each drug class in an unbiased manner, we performed PCA on the Z scored guide RNA counts per condition. PCA identified several important axes of variation, with PC1 primarily separating the DMSO from the treated samples, and PC2 distinguishing the molecular glues and classical inhibitors (**Figure S6**). Consistent with our previous findings, this analysis revealed that molecular glues enriched sgRNAs with PAMs targeting amino acids 75, whereas classical inhibitors enriched PAMs targeting amino acids 63 (**Figure S6b**). We note that sgRNA enrichment with CBE did not converge on a clear pattern, owing to known weaker efficacy of CBE and the lower coverage of the sgRNA library around the switch II pocket (**Figure S2a, S2b**).

**Figure 2.**
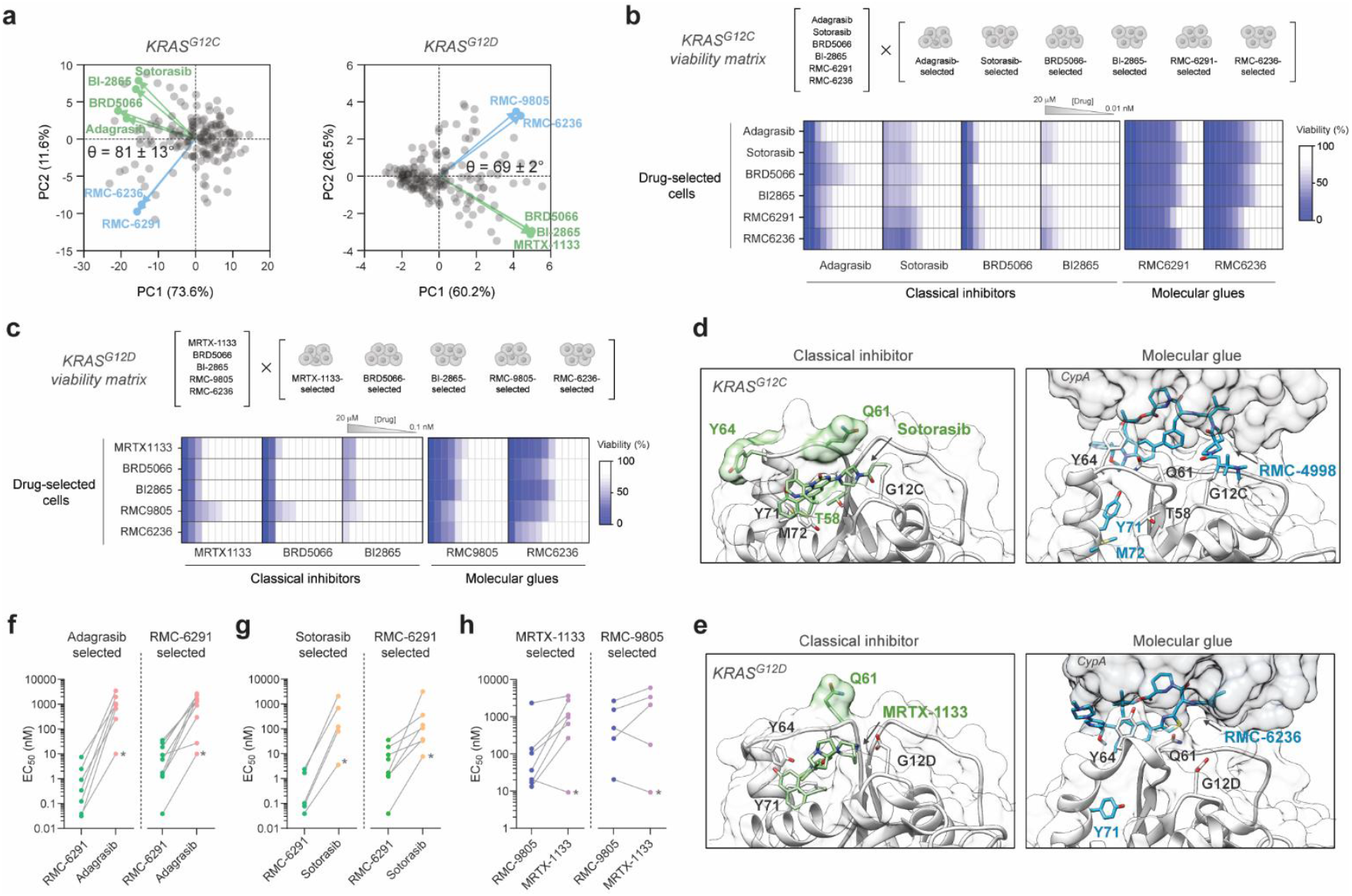
Molecular glues overcome resistance mutations against classical inhibitors. **a** Principal component analysis (PCA) biplot for sgRNA enrichment, using the drug selections at EC_80_ as variables. Lines correspond to loadings, highlighting the contribution of individual inhibitors. Respective average angles (θ) between the loadings for classical inhibitors (green) and molecular glues (blue) are indicated. Data for ABE transduced KRAS^G12C^ and KRAS^G12D^ selections are shown. **b–c** Viability matrix heatmaps showing inhibitor sensitivity of different selections in KRAS^G12C^ (**b**) and KRAS^G12D^ (**c**) cells transduced with ABE. Rows indicate cells from different drug selections, and columns represent drugs evaluated in the viability matrix assay. Each box of the heatmap shows the residual cell viability of a single dose across three independent replicates. Drugs were dosed with 20 μM maximal concentration and 3-fold dilution factor. **d–e** Structural mapping of the identified orthogonal resistance mutations in KRAS^G12C^ (**d**) or KRAS^G12D^ (**e**). Representative examples of classical inhibitors forming a binary complex (Sotorasib [PDB: 6IOM], MRTX-1133 [PDB: 7RPZ]) or molecular glues that form a ternary complex (RMC-4998 [PDB: 8G9P], RMC-6236 [PDB: 9AX6]) with CypA (dark gray) are provided. **f–h** Pair-wise comparison of adagrasib *versus* RMC-6291 (**f**), sotorasib *versus* RMC-6291 (**g**), and MRTX-1133 *versus* RMC-9805 (**h**) showing the drug sensitivity of individual sgRNA transduced cells. Nodes represent mean EC_50_ values of three independent replicates for cell viability measured following the same experimental outline as in panels **b** and **c.** EC_50_ values were interpolated by non-linear regression within the dose range of viability measurements. The drug indicated above each graph was used to select resistant cells. Drug sensitivity in the parental cell line is indicated by an asterisk in each graph.

Next, we performed a viability matrix assay in which we treated each resistant population with inhibitors that were not used for their respective selection. We observed orthogonal resistance between the two inhibitor classes, given that when classical inhibitors were used for selection, molecular glues did not display resistance, yet irrespective of the drug used for selection, classical inhibitors displayed resistance (**Figure 2b, 2c**). In all cases, molecular glues remained the most potent inhibitor class (**Figure 2b, 2c**). This result aligns with recent reports for patient-derived orthoxenografts, for which sotorasib resistance was effectively countered by RMC-6291.^30^

We next determined if the observed orthogonality in drug resistance also correlates with the underlying mutations (**Figure 1f, 1g** and **Figure S4**). Confirming our observation from the viability matrix assay, the molecular glues also showed orthogonal resistance mutations to classical inhibitors in KRAS^G12C^ cells, displaying lower incidence of T58, Q61, and Y64 mutations and higher incidence of Y71 and M72 mutations upon selection. Although Y71 and M72 do not directly interact with the molecular glues (**Figure 2d, 2e**), these compounds target the KRAS “on” state that bears a Y71-in conformation^31^, which could be disturbed by Y71 or M72 mutations. KRAS^G12D^ cells selected with molecular glues also showed enriched Y71 mutations, along with a higher incidence of Q61 and Y64 resistance mutations that overlap with classical inhibitors. While the Q61 mutation confers resistance against classical drugs by impairing GTPase activity and favoring the KRAS “on” state,^32^ it may also sterically hinder the binding of the RMC-9805 molecular glue. Based on AlphaFold3 modeling, Q61R and G12D are in close proximity and could potentially form a strong salt bridge, which renders G12D less available to interact with RMC-9805. In contrast, Y64 mutations could weaken critical contacts for binding to both classical inhibitors and molecular glues (**Figure 2d, 2e**). However, the activity of the molecular glues is likely less affected by Q61 or Y64 mutations, as they were still able to inhibit the growth of cells selected with classical inhibitors (**Figure 2c**). As observed in the sgRNA enrichment, deep sequencing for selections performed with CBE did not converge on a clear resistance pattern (**Figure S4a, S4b**).

To further confirm the orthogonality in resistance profiles of classical and molecular glue inhibitors, we performed validation studies by individual sgRNA transduction. sgRNAs with a >2 log_2_-fold enrichment were transduced along with corresponding base editors and subjected to respective drug selection for 4 weeks (**Figures S7, S8**). We then assessed the drug sensitivity of enriched cells to their respective selection drugs to confirm their resistance and subjected their genomic DNA to deep sequencing to identify resistance mutations (**Figures S9–S12**). Among the 66 sgRNA–drug pairings tested for KRAS^G12C^, 34 (52%) validated resistance.

Similarly, for KRAS^G12D^, 26 of 55 pairings (47%) showed enrichment of drug-resistant cells. Notably, most sgRNAs that did not show resistance were transduced with CBE and sgRNA pairings, which correlates with our results from the pooled screening. Upon performing matrixed viability studies in these individual sgRNA transductions, we further confirmed several key general orthogonalities between the two inhibitor classes (**Figure 2f–h** and **Figure S10**). First, regardless of the inhibitor type used for selection, a molecular glue consistently demonstrated greater potency than classical inhibitors. Second, with few exceptions, classical inhibitors exhibited more resistance (change in EC_50_) compared to molecular glue inhibitors, even for selections performed against molecular glues (**Figure 2f–h**). Finally, irrespective of the inhibitor type used for selection, nearly all classical inhibitors display resistance (**Figure S10**). We also confirmed the same mutations observed in the pooled screening by deep sequencing. Notably, we confirmed that the activity of pan-KRAS inhibitor RMC-6236 is more susceptible to mutations on the Y64 residue and less sensitive to Q61 mutations than the mutant-selective KRAS^G12C^ or KRAS^G12D^ inhibitors RMC-6291 and RMC-9805, respectively (**Figures S11, S12**). Although Y64 forms similar interactions with covalent and non-covalent RMC analogs^14, 16, 33^, the covalent nature of RMC-6291 and RMC-9805 may help to reduce the dependency on other interactions in the tertiary complex. This suggests that molecular glues can be designed to favor certain contacts within the ternary complex, a promising avenue to rapidly overcome emerging drug resistances. For example, a combined treatment of covalent and non-covalent molecular glues (e.g., RMC-9805 and RMC6236) could provide a therapeutic benefit by extending coverage against potential escape mutations.

In summary, we find that orthogonal resistance between inhibitors was observed irrespective of whether a chemical (cell pool selection) or genetic (sgRNA selection) perturbation was used to homogenize the cell population. Notably, molecular glue inhibitors overcame escape mutations of classical inhibitors and maintained cell growth inhibition at relevant concentrations <1 μM, but the opposite was not the case. We also identified escape mutations affecting drug response irrespective of the modality and found differences among the induced-proximity inhibitors. Our findings suggest that the development of molecular glues is not subject to the same restrictions as classical inhibitors on the target protein—the extended binding surface of the ternary complex provides an avidity effect by leveraging multiple contacts between recruited protein (CypA) and target (KRAS) and can be used to circumvent potential resistance mutations. While the development of molecular glues and other chemical inducers of proximity (CIP) remains challenging, emerging platforms such as CIP-DELs that screen large DNA-encoded libraries for simultaneous engagement of both a target and presenter protein can already enable rapid development of molecular glues targeting neo substrates.^34^ We anticipate that such platforms will synergize with the tiling mutagenesis and variant scanning strategies to prioritize the development of inhibitors with orthogonal resistance mechanisms.

## Experimental Methods

### Cell culture and chemical compounds

The following cell lines were obtained from ATCC: A549 (CCL-185), AGS (CRL-1739), MIA PaCa2 (CRL-1420), NCI-H1299 (CRL-5803), NCI-H727 (CRL-5815), and PSN-1 (CRL-3211). HEK293FT cells (Thermo R70007) were a kind gift from the F. Zhang lab and originally obtained from Thermo Fisher Scientific. All cells were cultured at 37°C and 5% CO_2_ in a humidified incubator and full growth media containing 10% (v/v) FBS (Gibco 16140-071), 100 units/mL penicillin and 100 μg/mL streptomycin (Gibco 15140-122). A549 and AGS cells were maintained in F12K media (ATCC 30-2004), MIA PaCa2 and HEK293FT cells were maintained in DMEM + GlutaMAX-I (Gibco 10564-011), and NCI-H1299, NCI-H727, and PSN1 cells were maintained in RPMI (Gibco 22400-089) medium. Chemical compounds were purchased from commercial sources (MedChemExpress) and used without further purification.

### Mapping residues interacting with inhibitors

X-ray structures of KRAS-inhibitor complex (PDB: 6OIM, 6UT0, 8AZY, 7RPZ, 8G9P, 8TBK, 9AX6) were loaded from the Protein Data Bank (rcsb.org), and prepared using Molecular Operating Environment (MOE 2022, Montreal, Canada). Preparation steps included adding hydrogen by Protonate3D, deleting water far from the binding site (> 4.5 Å), and fixing atoms more than 8 Å away from the ligand while preserving the coordinates of all heavy atoms. The interaction between inhibitors and KRAS was analyzed using both Ligand_Interactions and Contacts modules of MOE. A comparative ligand interaction fingerprint was also generated using the PLIF module. For BRD5066, the binding mode was generated using induced fit docking on PDB: 7RPZ. The docking was run with the Triangle Matcher algorithm (Amber10:EHT force field), London dG scores for placement and GBVI/WSA dG scores for refinement. The final figures were generated from PDB files or docking pose using UCSF Chimera 1.17.3. The AlphaFold3 predicted model was generated by inputting the KRAS4b sequence (residues 1-169) bearing G12D Q61R with GTP as a co-factor.

### Viability profiling of inhibitors in different KRAS dependent cell lines

Compound stocks were prepared at 10 mM concentration in DMSO solvent. Compounds were plated into white 384-well plates at maximal concentration of 20 μM and diluted into a 12-point dose response with a 3-fold dilution factor using a Tecan D300e compound dispenser. All wells were normalized to 0.5% (v/v) DMSO content. Cells were plated at a density of 1,000 cells/well in 25 μL full media (10% (v/v) FBS and 1% (v/v) penicillin/streptomycin). The plates were incubated in a 37°C, 5% CO_2_ humidified incubator for 72 h. A cell Titer Glo (Promega) assay was performed by adding a reagent volume equal to half the total volume of the well. Plates were read on an Envision plate reader. Percentage viability was calculated by normalizing to DMSO only controls. Data were fitted to a four-parameter logistic curve for dose-response inhibition using GraphPad Prism 10.

### sgRNA library construction for base editing

The base editor library was constructed by adapting previously established protocols^23, 35^. A custom library of sgRNA sequences tiling across the KRAS4b isoform was designed using the CRISPOR tool against the GRCh38/hg38 genome assembly^26^. Genomic DNA sequences corresponding to the four exons of KRAS were used and sgRNAs with specificity scores below 20 were excluded. Aside from 20 bp flanking the exons, intronic regions were omitted for designing the library. Control non-targeting sgRNAs that induce neutral effects on gene editing (i.e., targeting safe harbor loci or intergenic regions) were included as well and the library was ordered with corresponding homology arms for Gibson assembly as an oligonucleotide pool from Twist Bioscience (a detailed list of all sgRNAs and their sequences is provided in **Data S1**). Sanger Sequencing (Genewiz) was used to analyze the genomic sequence of the KRAS4b isoform in MIA PaCa-2 and AGS cells to ensure library coverage (primer list and exon sequences are provided in **Table S1**). The sgRNA plasmid library was constructed using Gibson assembly. In brief, the oligonucleotide pool was PCR amplified and purified with a QIAquick PCR purification kit (Qiagen, 28104). ABE8e (a gift from John Doench & David Root, Addgene 179099)^28^ and BE3.9 (a gift from John Doench & David Root, Addgene 179096)^28^ vectors were linearized with FastDigest Esp3I (ThermoFisher FD0454) and FastAP Thermosensitive Alkaline Phosphatase (Thermo EF0654) in buffer supplemented with DTT. Gibson assembly was used to construct the plasmid library with NEBuilder HiFi DNA Assembly Master Mix (New England Bio Labs E2621L) following the manufacturer’s instructions. The pooled plasmid library was isolated by isopropanol precipitation and electroporated into Endura DUO electrocompetent cells (Biosearch Technologies 60242) using a Gene Pulser Xcell Electroporation System (Bio-rad). The cells were plated on LB plates containing 100 μg/ml ampicillin, and transformation efficiency was confirmed to be > 10’000 times the number of sgRNAs in the library. Cells were harvested and plasmids isolated using ZymoPure II Plasmid Maxiprep Kit (Zymo Research D4203) and purified from endotoxins.

### NGS and analysis of library coverage

Primers annealing on the 3’ and 5’ end of the protospacer sequence were to PCR amplify the sgRNA barcodes and attach common adapters for Illumina sequencing (22 cycles were performed using NEBNext Ultra II Q5 Master Mix [New England Bio Labs M0544L]). PCR products were purified using SPRIselect beads (Beckmann Coulter Life Sciences B23317), following the manufacturer’s instructions to elute approximately 300–500 bp oligonucleotide fragments. The library was diluted to 2 nM using a KAPA Library Quantification Kit (Roche 07960140001) on a QuantStudio 7 Flex qPCR instrument. Sequencing was performed on an Illumina MiSeq instrument using a Micro Kit v2 (Illumina MS-103-1002) or Nano Kit v2 (Illumina MS-103-1001) following the manufacturer’s instructions. The NGS data was analyzed using previously described Python scripts to measure the sgRNA distribution.^35^ Briefly, the number of sequencing reads that exactly match a protospacer sequence was counted and converted to reads per million. Sequences without any positive match were increased by a pseudo count of 1, and all counts were log_2_-transformed. The resulting values were used to calculate the sgRNA library distribution and enrichment.

### Lentiviral transduction of sgRNA library

18 × 10^6^ HEK293FT cells were seeded in a 15-cm dish and left to adhere overnight at 37°C in a humidified 5% CO_2_ incubator. Upon reaching approximately 90% confluency, the cells were transfected with lentiviral plasmids. 1.2 mL Opti-MEM (Gibco 31985062), 3 μg pMD2.G plasmid (a gift from Didier Trono, Addgene 12259), 6 μg psPAX2 (a gift from Didier Trono, Addgene 12260), 9 μg sgRNA plasmid library, and 60 μL FuGENE HD transfection reagent (Promega E2311) were gently mixed in an Eppendorf, incubated 10 min at room temperature, and slowly added directly to the cell media for transfection. Cells were moved back to the incubator for 6 hours, and media was replaced with fresh complete DMEM + GlutaMAX-I (Gibco 10564-011). The cells were placed back into the incubator for another 48 hours before collecting the lentiviral particles by filtering the cell media through a 0.45 μm filter. Supernatant was collected and lentiviral titer estimated using Lenti-X GoStix Plus (TaKaRa Bio 631280). To transduce the lentivirus, AGS (KRAS^G12D^) or MIA PaCa2 (KRAS^G12C^) cells were seeded in 12-well plates at a density of 1.5 × 10^6^ cells per well in the presence of 8 μg/mL polybrene (Santa Cruz Biotechnology). Transduction was performed by spinfection at 1,800 × g at 37°C for 90 minutes, adding the appropriate amount of lentivirus to obtain a multiplicity of infection (MOI) < 0.3. After spinfection, the cells were gently resuspended and diluted with fresh media. 48 hours post-transduction, transduced cells were selected for at least 7 days using 2 μg/mL puromycin (Gibco A1113803). Genomic DNA was extracted from the selected cells using a QIAamp DNA Blood Mini Kit (Qiagen, 51104) according to the manufacturer’s instructions. The region corresponding to the sgRNA was PCR-amplified to incorporate Illumina adapters, and the coverage and distribution of the sgRNA library were assessed by NGS as described above for the plasmid library.

### Selection and sequencing of drug-resistant cells

AGS (KRAS^G12D^) and MIA PaCa2 (KRAS^G12C^) cells transduced with either the base editor and the sgRNA library were split into three replicates for each drug or vehicle selection. Cells were initially treated with the lowest drug concentration (EC_30_, relative to parental cell line sensitivity). After two weeks, the dose was increased to an intermediate concentration (EC_50_) and further escalated to the highest concentration (EC_80-90_) after another two weeks. A detailed dosing regimen is provided in **Table S2**. 0.1% DMSO was maintained throughout the selection. Following six weeks of drug treatment, cells were collected for drug sensitivity measurements, sgRNA distribution analysis, and genotyping. Genomic DNA was extracted using the QIAamp DNA Blood Mini Kit (Qiagen, 51104), and sgRNA distribution was assessed as described for the plasmid library. sgRNA distribution after drug selection was normalized to the distribution immediately after lentiviral transduction (also referred to as “plasmid library”) to calculate a resistance score. sgRNAs with a resistance score (i.e., a log_2_-fold change) greater than 2 compared to the plasmid library were considered enriched and selected for validation. This threshold corresponds to an average enrichment of more than two standard deviations above the mean of non-targeting sgRNA controls across all drug selections. Detailed sgRNA resistance scores for each drug selection after two, four, and six weeks of treatment are provided in **Data S2–S3**.

### Statistical Analysis

Principal component analysis was performed by GraphPad Prism 10. Angles between loadings were quantified by angle analysis in Fiji, which measured angle difference between each loading and the abscissa in the indicated biplot. Spearman correlation was computed using GraphPad Prism 10 with a two-tailed P value and 95% confidence interval. To perform the unbiased principal component analysis, normalized guide counts were converted to a Z score using the distribution of each guide in the DMSO condition as the background. Then, to minimize the signal of unaffected guides, guides that did not display a Z score of 3 or greater in at least 1 condition were filtered. Finally, PCA and loadings were calculated and visualized using the functions available in the Scanpy package (https://scanpy.readthedocs.io/en/stable/). Statistical significance comparing drug-selected lines to parental was determined by one-way ANOVA (* p < 0.05, ** p < 0.01, *** p < 0.001, **** p < 0.0001) using GraphPad Prism 10.

### Viability Matrix Assay

Compound stocks were prepared at 10 mM concentration in DMSO solvent. Compounds were plated into white 384-well plates at maximal concentration of 20 μM and diluted into a 12-point (AGS) or 14-point (MIA PaCa2) dose response with 3-fold dilution factor using a Tecan D300e compound dispenser. All wells were normalized to 0.5% (v/v) DMSO content. Cells were plated at a density of 500 cells/well in 25 μL full media. The plates were incubated at 37°C, 5% CO_2_ humidified incubator for 72 h. A Cell Titer Glo (Promega) assay was performed by adding reagent volume equal to half the total volume of the well. Plates were read on an Envision plate reader. Percentage viability was calculated by normalizing to DMSO-only controls. Data were fitted to a four-parameter logistic curve for dose-response inhibition using GraphPad Prism 10.

### Genotyping

Genomic DNA was isolated from drug-selected cells using the QIAamp DNA Blood Mini Kit (Qiagen, 51104), as previously described for sgRNA distribution analysis. Since KRAS exons are within the 500-base-pair limit for paired-end Illumina MiSeq sequencing, amplicons corresponding to the full exons of the KRAS4b isoform were sequenced for each drug selection and joined for the analysis. At least 1.5 μg of DNA was used as a template for the first PCR round, generating gene-specific amplicons with common overhangs (18–20 cycles, NEBNext Ultra II Q5 Master Mix, New England Bio Labs M0544L). The PCR products were then diluted 1:10 and subjected to a second PCR round with primers containing barcoded Illumina adapters (15 cycles, NEBNext Ultra II Q5 Master Mix). Details for the primer sequences and the indexes used for NGS multiplexing are provided in **Table S3**. SPRIselect beads were used to isolate the PCR products. The library was diluted to 2 nM using a KAPA Library Quantification Kit (Roche 07960140001) using a QuantStudio 7 Flex qPCR instrument and sequenced on an Illumina MiSeq instrument using a v2 Kit (Illumina MS-103-1003 or MS-102-2003) following the manufacturer’s instructions. Reads were aligned to their reference amplicons using CRISPResso2 (ref^36^), and the resulting alleles analyzed via a custom Python script. In brief, reads were trimmed to the coding sequence of KRAS4b, allele frequency data were translated to the amino acid level, and substitution frequency data were consolidated from across reads. Mutational frequency was then determined by summing the frequencies of all alleles carrying a mutation at the same site (e.g., Q61). Hotspot mutations were defined as sites with a mutational frequency greater than 10%. Allele deep sequencing and mutational frequency data are provided in Supplementary Data 4 and 5 for the base editor screening and sgRNA validation, respectively.

## Delivery of sgRNAs for validation

ABE (Addgene 179099) and CBE (Addgene 179096) plasmids were digested with FastDigest Esp3I (ThermoFisher FD0454) and FastAP Thermosensitive Alkaline Phosphatase (Thermo EF0654) in the presence of DTT. The sgRNA oligonucleotides were PCR amplified using NEBNext Ultra II Q5 Master Mix (New England Bio Labs M0544L) and individually subjected to Gibson assembly using NEBuilder HiFi DNA Assembly Master Mix (New England Bio Labs E2621L). A detailed list of all oligonucleotides used for sgRNA validation is provided in **Table S4**. The products were transformed into competent cells (New England Bio Labs C2984H) and propagated in LB supplemented with 100 μg/mL ampicillin with shaking at 30°C overnight. Plasmid was harvested and purified using the ZymoPURE II Plasmid Midiprep Kit. Plasmid sequence was verified through sequencing by Plasmidsaurus. The purified plasmids were co-transfected with psPAX2 (Addgene 12260) and pMD2.G (Addgene 12259) lentiviral packing and envelope-expressing plasmids into HEK293FT cells using FuGENE HD (Promega). The media was replaced after 6 h post-transfection. 48 h later, the viral supernatant was collected, filtered (0.45 μm), and frozen at −80°C. To transduce the lentivirus, AGS or MIA PaCa2 cells were seeded in 12-well plates at 1.5 × 10^6^ cells/well and 8 μg/mL polybrene (Santa Cruz Biotechnology), lentivirus was titrated, and spinfection at 1800 × *g* at 37°C for 90 min was conducted. Following spinfection, the cells were gently resuspended, diluted with fresh media, and transferred to 6-well plates. 48 h post-transduction, the cells were selected with 2 μg/mL puromycin. Viral titer that yielded an MOI of ~0.3 was further selected and used for further experiments. Following 2 weeks of puromycin selection, cells were subjected to drug selection by treating at the EC_80_ concentration of the respective drug (0.1% (v/v) DMSO composition) for 4 weeks. Daily brightfield images of each flask and trypan blue exclusion counts of viable cells during each passage were acquired to monitor confluency and cell count, respectively, throughout the selection period. Confluency was measured by pixel classification and segmentation using Ilastik through features analysis of edges and texture. Segmented images were quantified by thresholding and area density measurements in Fiji.

## Acknowledgements

We thank B. Liau, H.S. Kwok, and A. Waterbury at Harvard University (Cambridge, MA) for insightful discussions. This work was supported by NIH (R21AI154099, R01GM137606, R01DK129464, R01GM132825). M.M was supported by the Swiss National Science Foundation (P500PN_214260), and KK acknowledges support from the Natural Sciences and Engineering Research Council of Canada Post-doctoral Fellowship.

## Notes

No conflict of interest

