## Supporting Information for "Orthogonal resistance mechanisms of classical- and induced-proximity inhibitors"

### Table of contents

#### Supplementary Figures

**Figure S1.** Current landscape of KRAS inhibitors. 3

**Figure S2.** Coverage of the sgRNA library. 4

**Figure S3.** KRAS variant scanning with adenine (ABE) and cytosine (CBE) base editors. 5

**Figure S4.** Deep sequencing reveals drug-resistant alleles. 6

**Figure S5.** Base editor variant scanning shows distinct correlations for classical inhibitors and molecular glues. 7

**Figure S6.** PCA forms distinct sgRNA enrichment clusters for classical inhibitors and molecular glues. 8

**Figure S7.** Growth curves for enrichment of $KRAS^{G12C}$ drug-resistant cells. 9

**Figure S8.** Growth curves for enrichment of $KRAS^{G12D}$ drug-resistant cells. 12

**Figure S9.** Delivery of individual sgRNA/BE pairing yields KRAS-resistant cells. 14

**Figure S10**. Viability-matrix with validated sgRNAs confirms drug-resistant KRAS variants. 15

**Figure S11.** Genotyping of $KRAS^{G12C}$ drug-selected cells transduced with validated sgRNAs. 16

**Figure S12.** Genotyping of $KRAS^{G12D}$ drug-selected cells transduced with validated sgRNAs. 17

#### Supplementary Tables

**Table S1.** Sanger sequencing of KRAS4b exons. 18

**Table S2.** Drug dosing regimen for base editor screening. 19

**Table S3.** Primers used for deep sequencing of KRAS4b exons. 20

**Table S4.** Oligonucleotides for validating sgRNAs. 21

#### Notes for Supplementary Data

**Data S1 Notes.** KRAS sgRNA library sequences. 23

**Data S2 Notes.** Base editor screening data in MIA PaCa2 ($KRAS^{G12C}$) cells. 23

**Data S3 Notes.** Base editor screening data in AGS ($KRAS^{G12D}$) cells. 23

**Data S4 Notes.** Deep sequencing of editing outcomes in base editor screening. 23

**Data S5 Notes.** Deep sequencing of editing outcomes for validated sgRNAs. 23

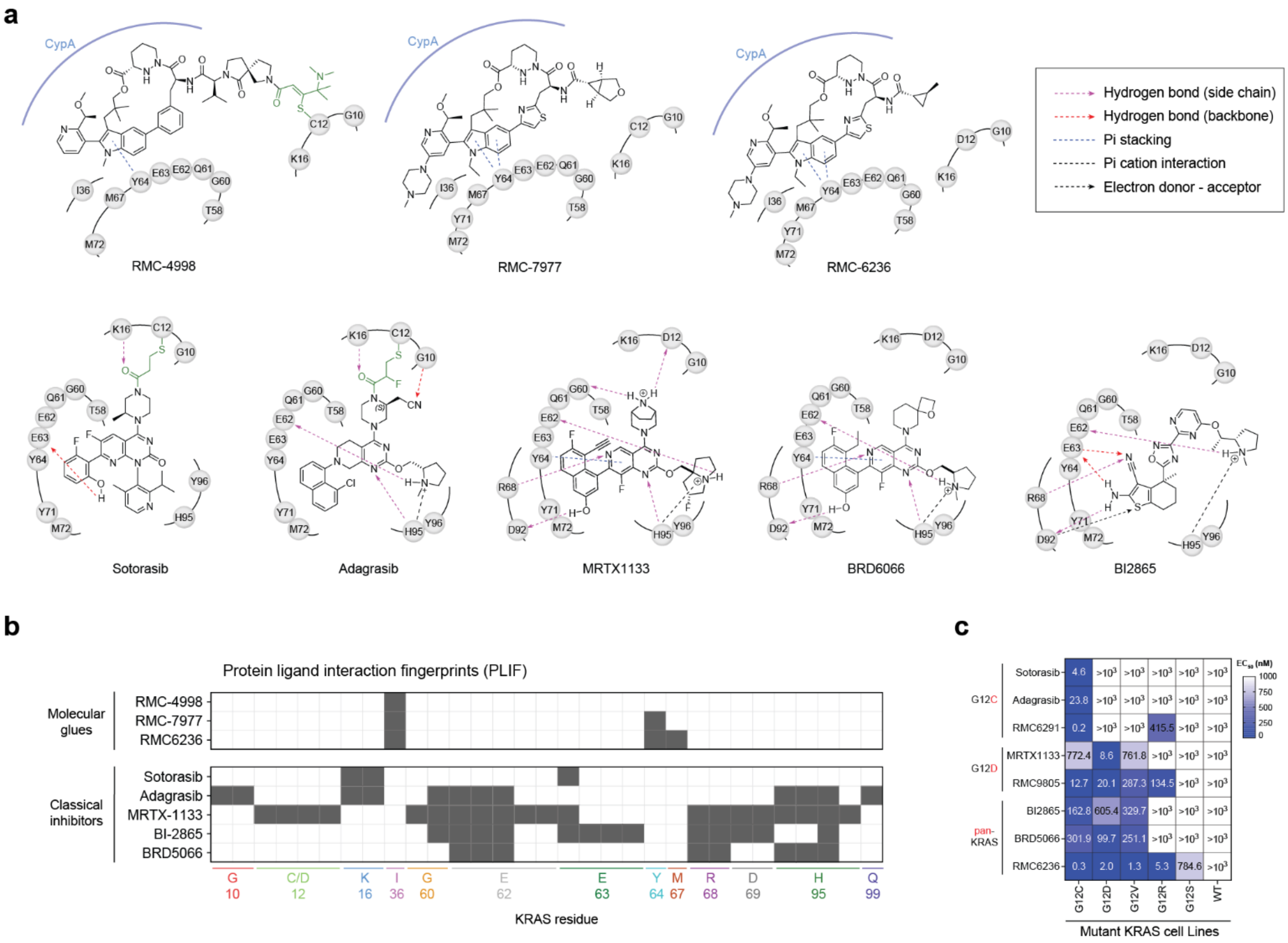


**Figure S1. Current landscape of KRAS inhibitors. a** The structures of different inhibitors and their molecular contacts with KRAS are shown. Most of them form multiple interactions with residues on switch II. These 2D representations were constructed from analyzing PDB files of the corresponding inhibitors with KRAS; RMC-4998 (8G9P), RMC-7977 (8TBK), RMC-6236 (9AX6), sotorasib (6OIM), adagrasib (6UT0), MRTX1133 (7RPZ), BRD6066 (docking model based on PDB:7RPZ) and BI2865 (8AZY); using Maestro (Schrodinger Suite) and MOE (Molecular Operating Environment). Hydrophobic interactions (other than pi-stacking) were omitted for clarity. **b** Protein Ligand Interaction Fingerprint (PLIF) of KRAS inhibitors are shown. The ligand interaction profile for each amino acid residue of KRAS is represented in a binary format using the default cutoff and energy estimation from MOE. For each interaction type (H-bond donor, H-bond acceptor, ionic attraction, metal attraction, arene attraction), the absence of interaction or very weak interaction < 0.5 kcal/mol was encoded as 00 (white box), noticeable interaction > 0.5 kcal/mol was encoded as 01 (one black box), and strong interaction > 1.5 – 3.5 kcal/mol was encoded as 11 (two black barcodes). Since a single residue can form multiple interaction types with a ligand (e.g., E62, E63), the map displays a summation of all interaction types. **c** Heatmap showing the efficacy of different inhibitors against mutant KRAS cell lines: G12C (MIA PaCa2) G12D (AGS), G12V (NCI-H727), G12R (PSN-1), G12S (A549), and WT (NCI-H1299). NCI-H1299 cell lines carry NRAS Q61K mutations and do not depend on KRAS inhibition. $EC_{50}$ of three independent replicates are shown.

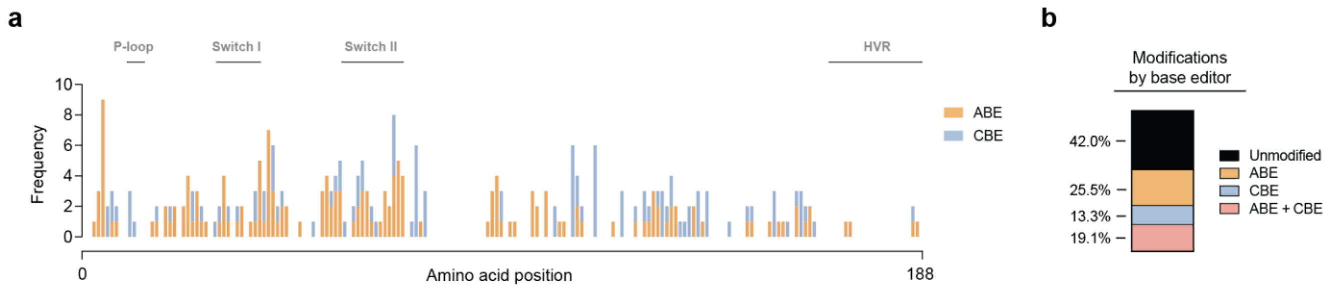


**Figure S2. Coverage of the sgRNA library. a** Number of predicted missense mutations for each base editor and calculated cumulative frequencies based on sgRNA mapping. Any C (for CBE) or A (for ABE) within the editing window +4 to +8 of the protospacer sequence was assumed to get converted to T or G, respectively. For each amino acid residue, the cumulative mutational frequency across all sgRNAs in the library was calculated for each base editor. Edited sites for each sgRNA and editor are provided in Data S1. **b** Relative number of modifications induced by either base editor. Residues that can get mutated at least once were counted. 58% of residues in KRAS can get mutated at least once.

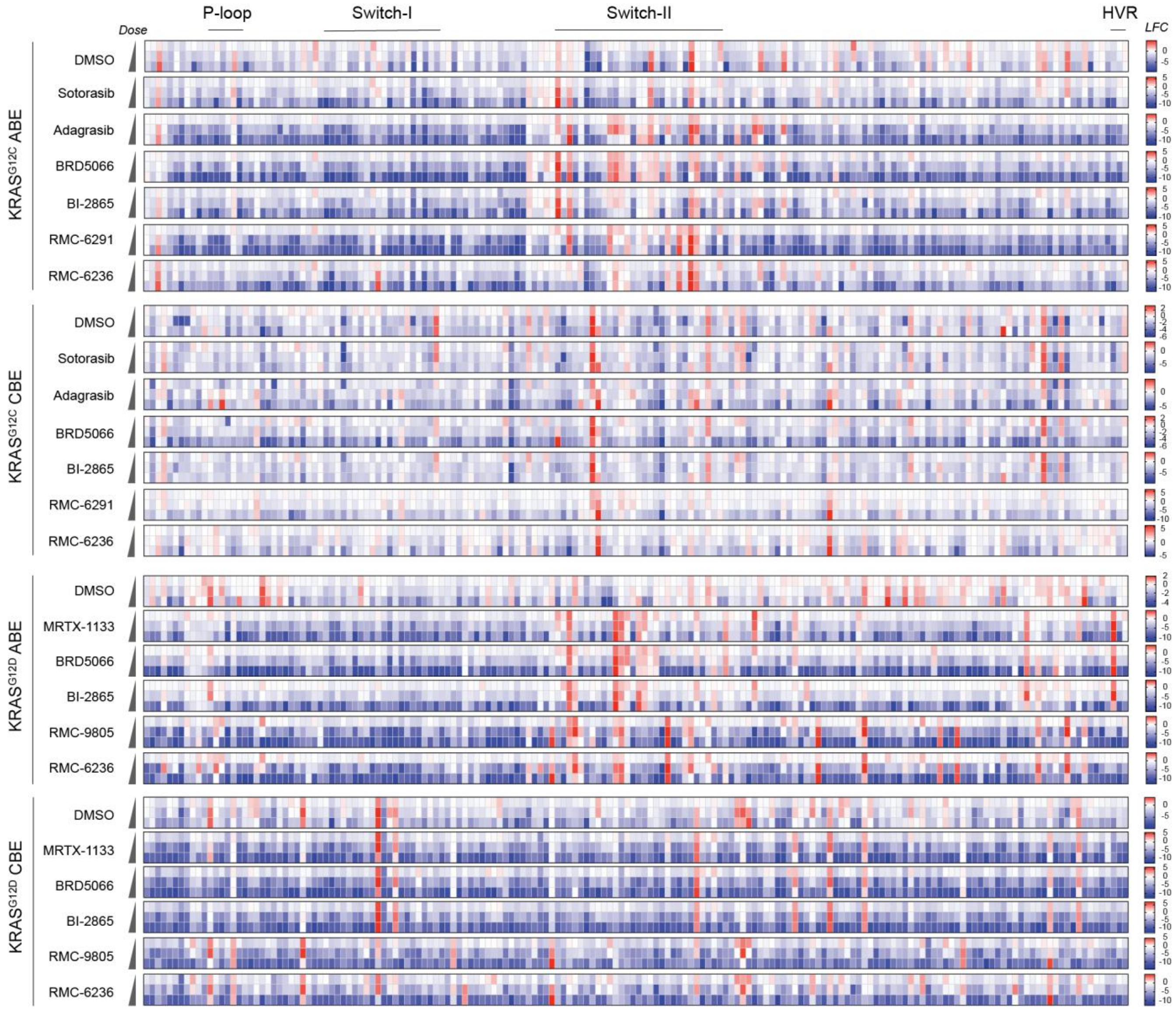


**Figure S3. KRAS variant scanning with adenine (ABE) and cytosine (CBE) base editors.** sgRNA enrichment expressed as $\log_2$ fold-change (LFC) versus the plasmid library across all drug treatments, drug concentrations, editors, and cell lines. Each data point in the heatmap represents the enrichment value of an sgRNA across three independent selection replicates. Detailed information of the dose regimen is provided in Table S2, and sgRNA resistance scores are provided in Data S2 and S3.

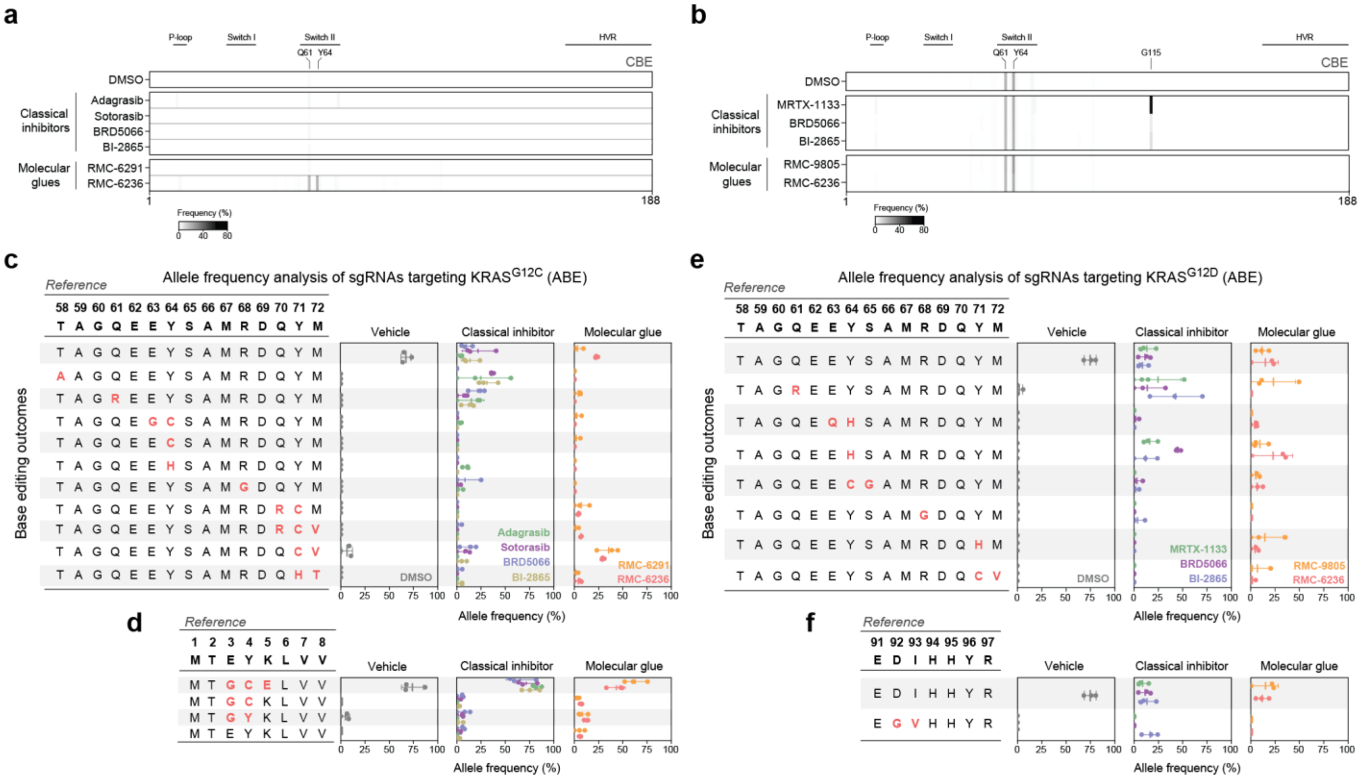


**Figure S4. Deep sequencing reveals drug-resistant alleles. a–b** Mutational frequencies at different sites in CBE edited KRAS$^{G12C}$ (**a**) and KRAS$^{G12D}$ (**b**) cells after six weeks of drug selection. Genomic DNA was extracted, and the four exonic regions of the *KRAS4b* isoform were PCR-amplified and subjected to deep sequencing. Mutational frequency was calculated as the sum of all alleles carrying a missense mutation at the indicated site. Hotspots with >10% modification are annotated on the heatmap. Frequencies represent the mean of three independent drug selections. **c–d** Allele frequencies in base-edited KRAS$^{G12C}$ cells after six weeks of drug selection. Each row represents an allele, with mutated sites highlighted in red. The reference sequence is shown at the top. Alleles with at least 10% frequency in any sample are displayed. Identical mutational profiles in the indicated regions were merged. Allele frequencies for vehicle-selected cells and cells treated with either a classical inhibitor or a molecular glue are shown on the right. **c** shows allele frequencies for residues T58-M72, while **d** shows frequencies for residues M1-V8. Mean and standard deviation are based on three independent selections. **e–f** Same as **c–d**, but for KRAS$^{G12D}$ drug selections. **e** represents allele frequencies for residues T58-M72, while **f** represents frequencies for residues E91–R97.

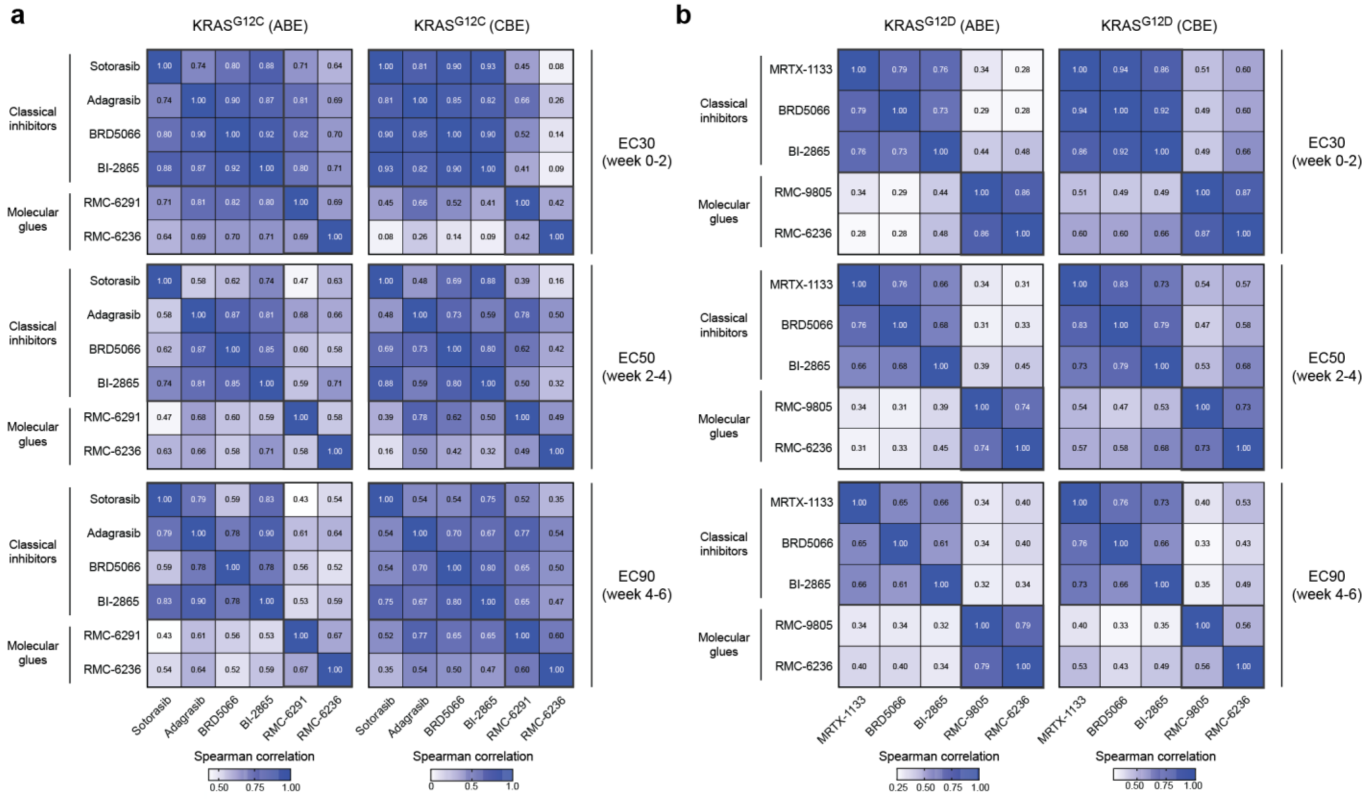


**Figure S5. Base editor variant scanning shows distinct correlations for classical inhibitors and molecular glues. a-b** Spearman correlation between sgRNA enrichment profiles in KRAS$^{G12C}$ (**a**) and KRAS$^{G12D}$ cells (**b**). Correlation was calculated using the mean of three independent replicates for each variant scanning experiment.

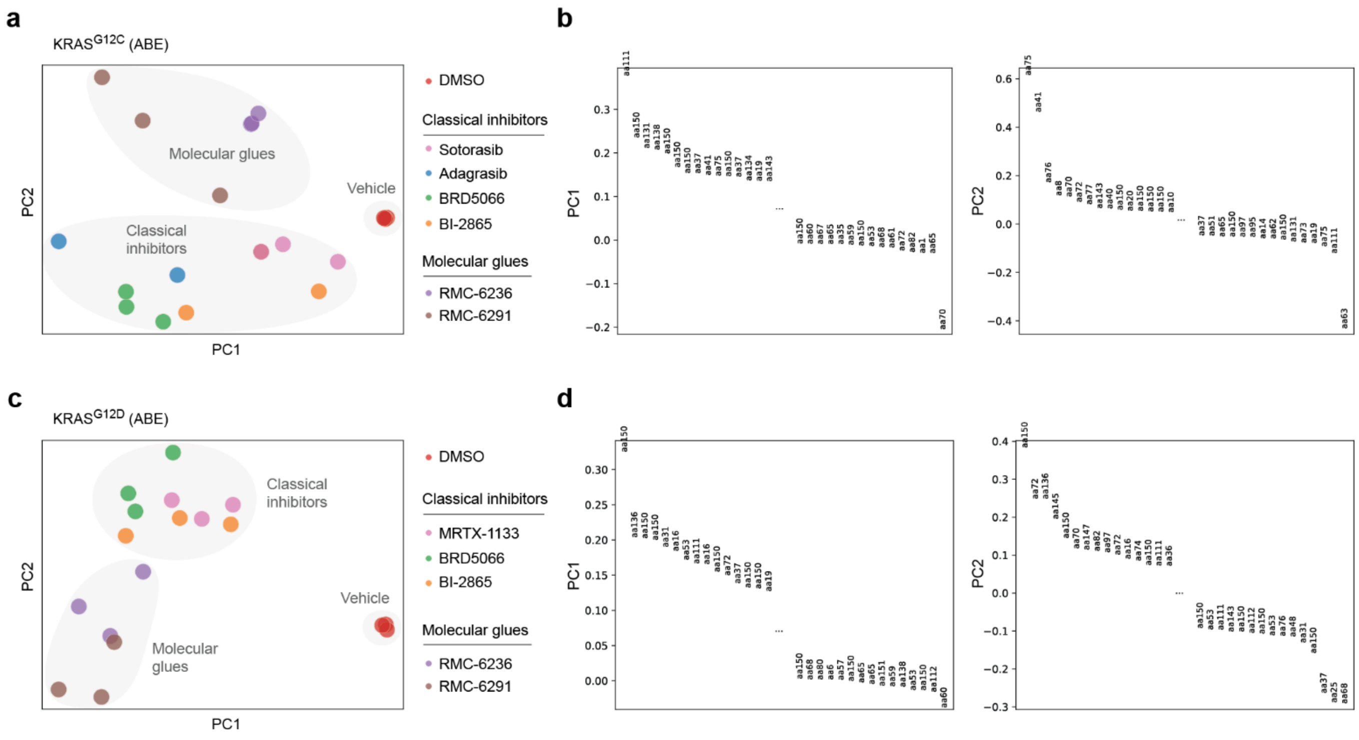


**Figure S6. PCA forms distinct sgRNA enrichment clusters for classical inhibitors and molecular glues. a–b** PCA (**a**) and loadings (**b**) for sgRNA enrichment profiles in $KRAS^{G12C}$ cells. **c–d** Same as **a** and **b**, but for $KRAS^{G12D}$ cells. Each datapoint in **a** and **c** represents a selection replicate. Loadings in **b** and **d** represent different sgRNAs and their targeted amino acid position relative to the PAM.

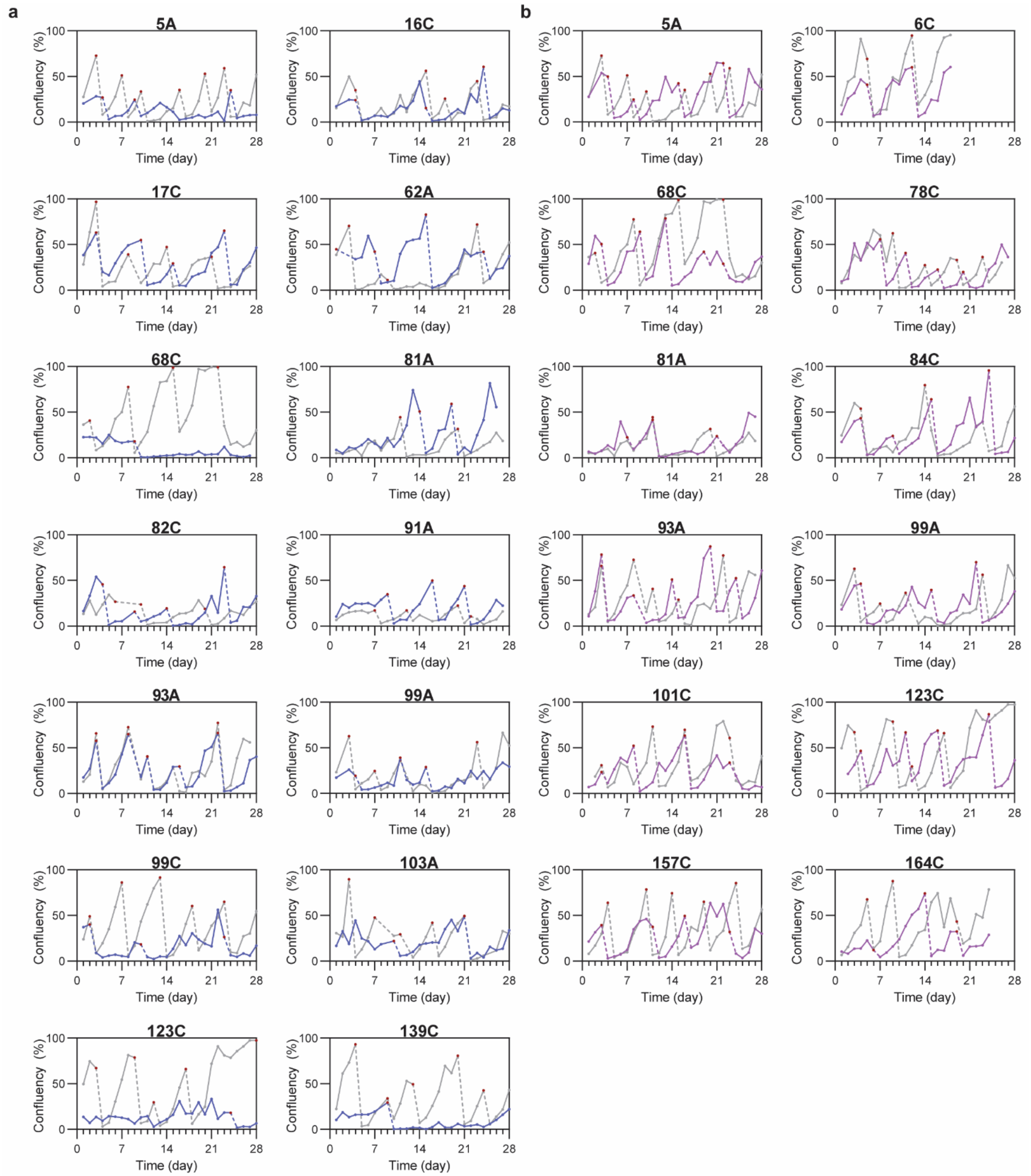


**Figure S7. Growth curves for enrichment of KRAS$^{G12C}$ drug-resistant cells.** Cell confluency (%) measured by brightfield microscopy plotted against time since start of drug selection with adagrasib (**a,** dark blue), sotorasib (**b**, purple), BI2865 (**c,** lavender), BRD5066 (**d,** teal), RMC6291 (**e,** green), or RMC6236 (**f**, pink) in cells individually transduced with indicated sgRNA (top of each panel). Red points and dashed lines indicate passage points. DMSO control selections are shown in grey traces and drug selections are shown in the indicated colors.

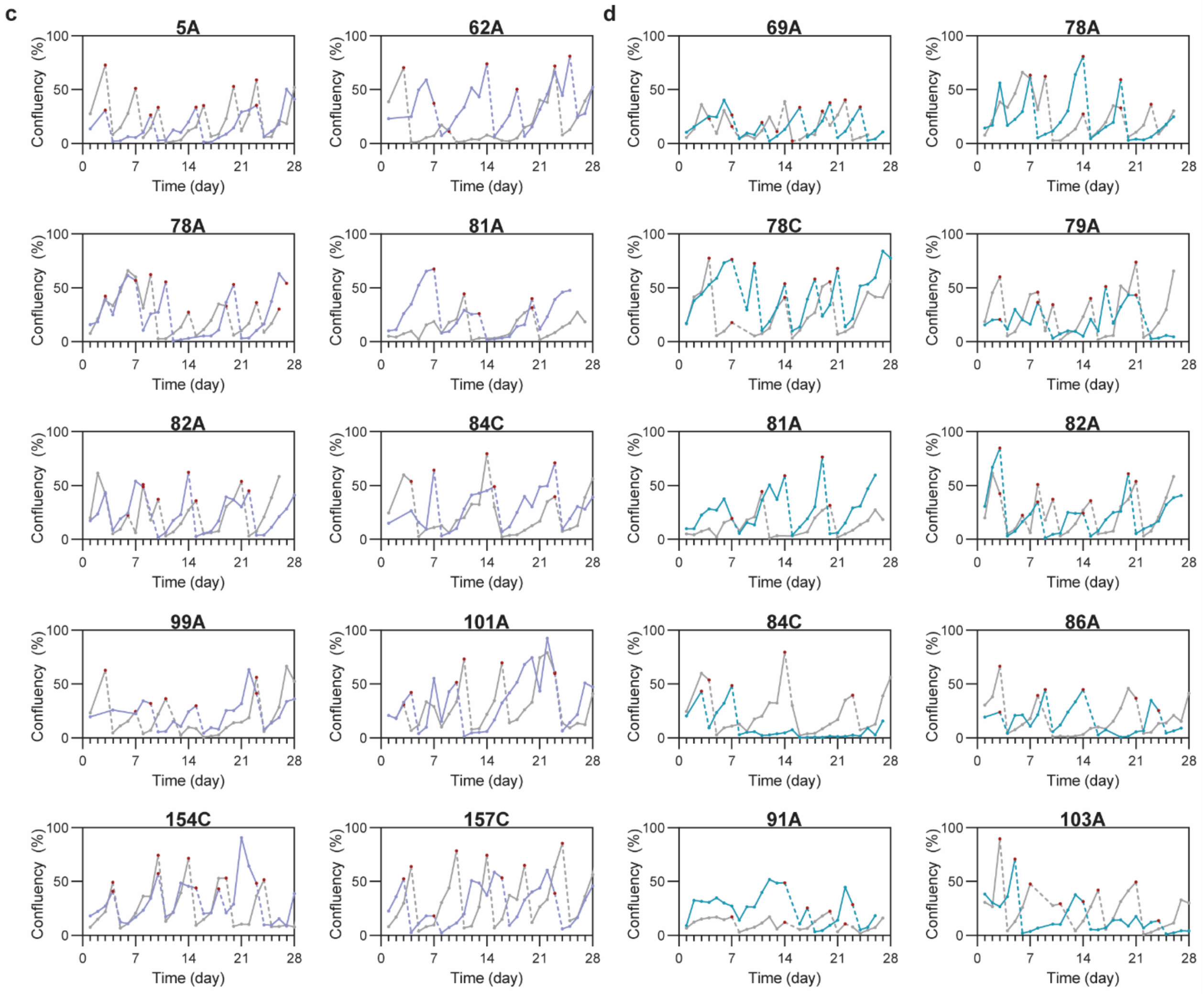


**Figure S7. Growth curves for enrichment of KRAS$^{G12C}$ drug-resistant cells.** Continued.

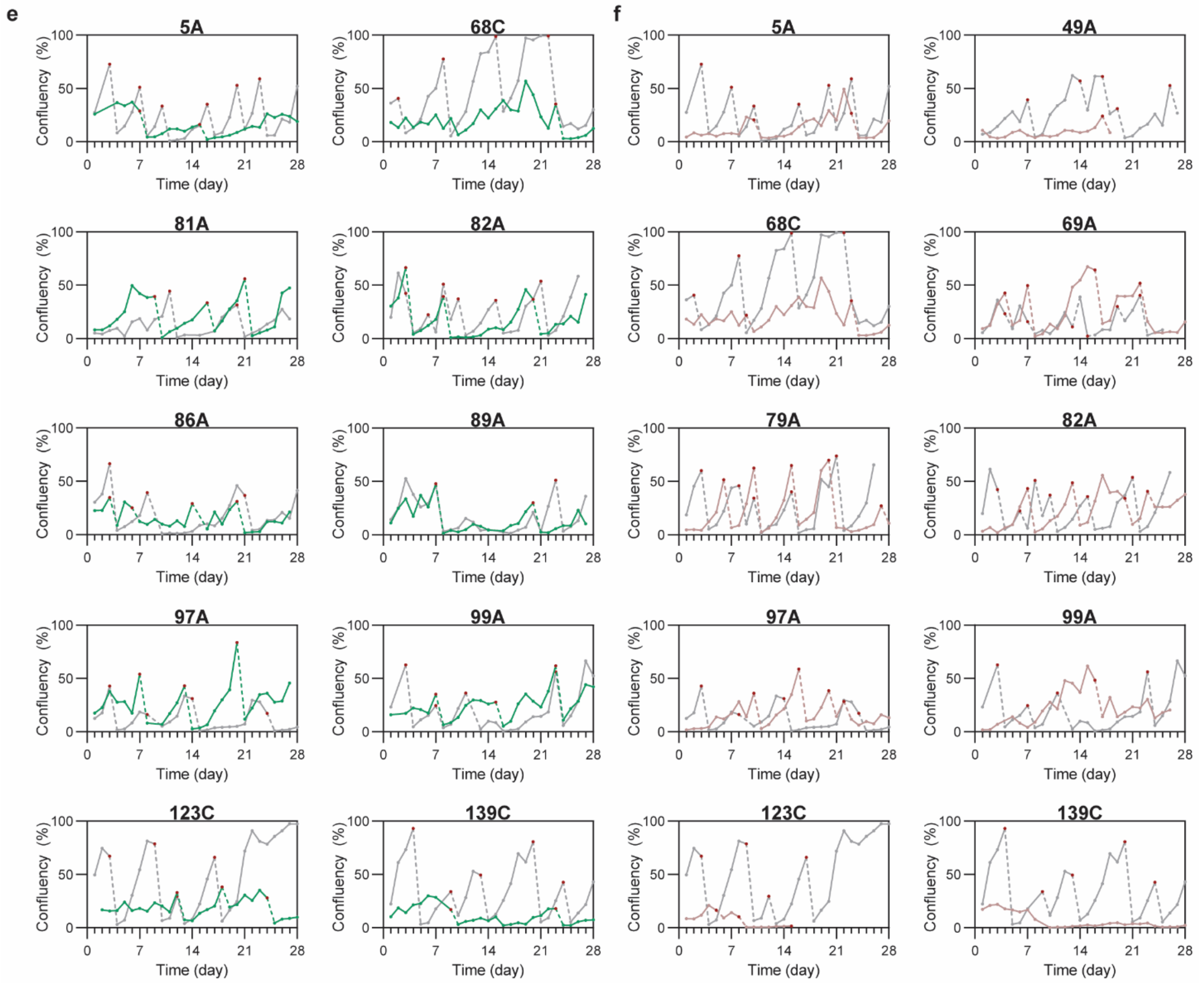


**Figure S7. Growth curves for enrichment of $KRAS^{G12C}$ drug-resistant cells.** Continued.

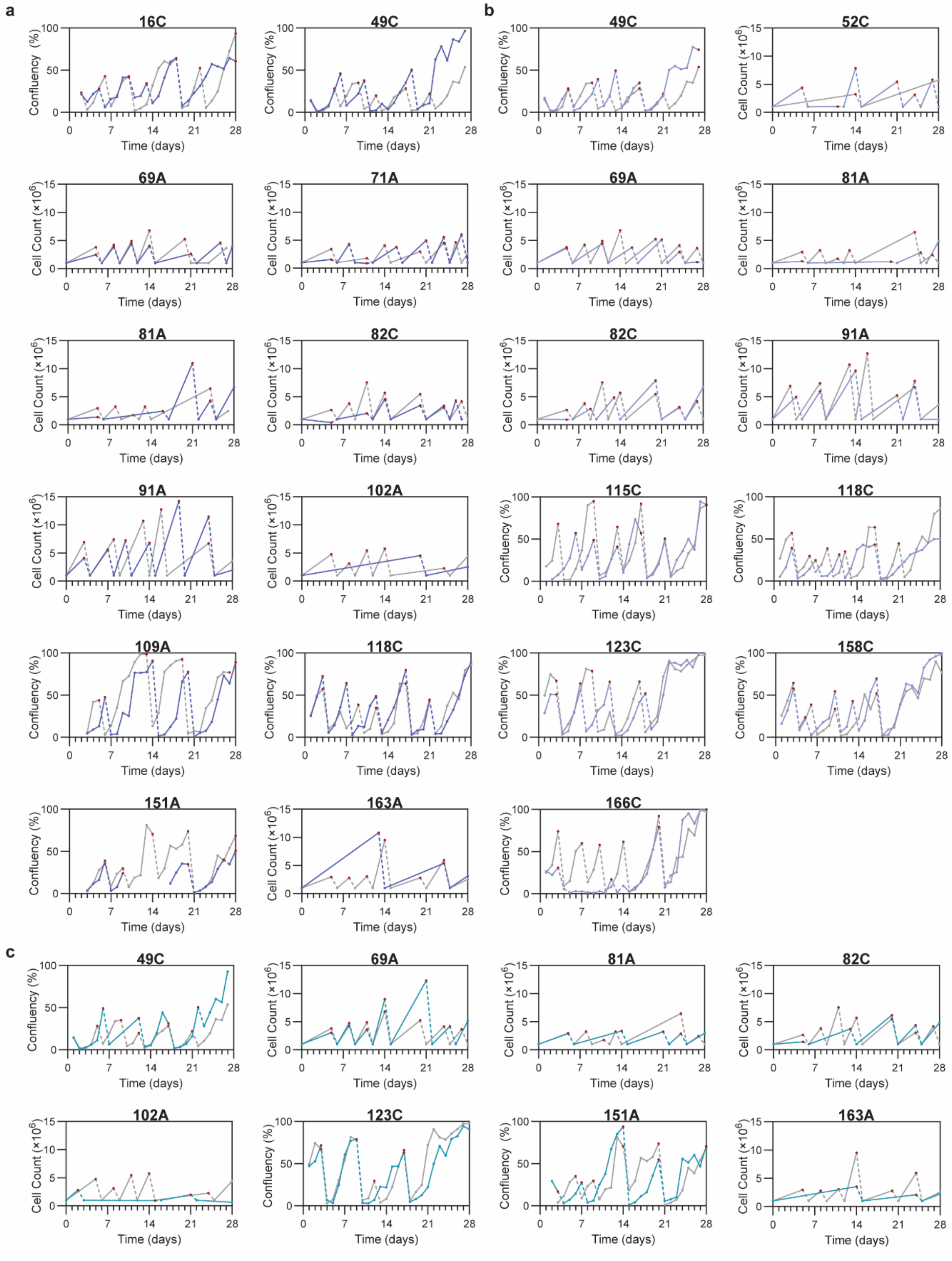


**Figure S8. Growth curves for enrichment of KRAS$^{G12D}$ drug-resistant cells.** Cell confluency (%) measured by brightfield microscopy or cell count measured by trypan blue counting plotted against time since start of drug selection with MRTX1133 (**a,** dark blue), BI2865 (**b,** lavender), BRD5066 (**c,** teal), RMC9805 (**d,** green), or RMC6236 (**e**, pink) in cells individually transduced with indicated sgRNA (top of each panel). Red points and dashed lines indicate passage points. DMSO control selections are shown in grey traces and drug selections are shown in the indicated colors.

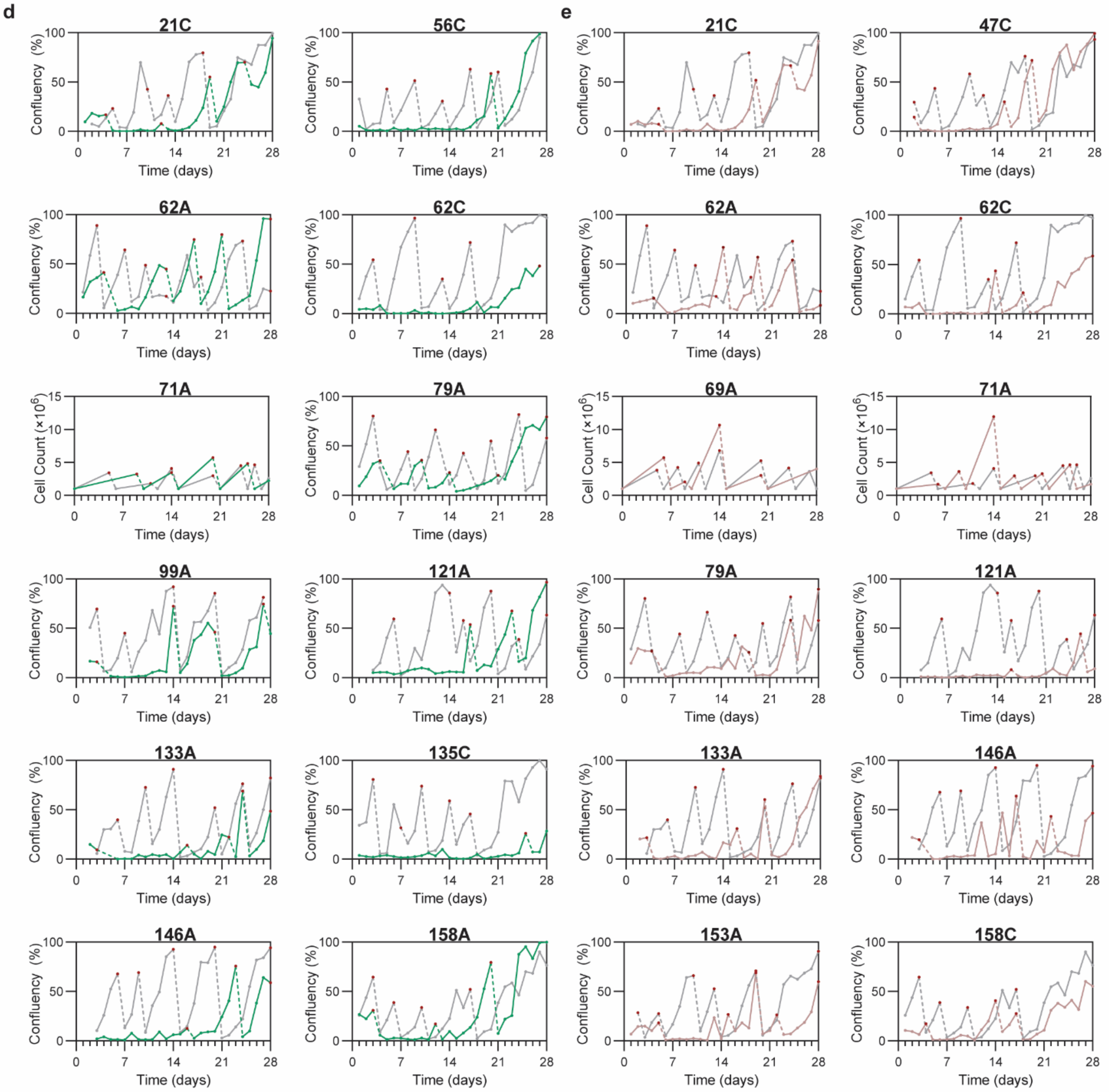


**Figure S8. Growth curves for enrichment of KRAS$^{G12D}$ drug-resistant cells.** Continued.

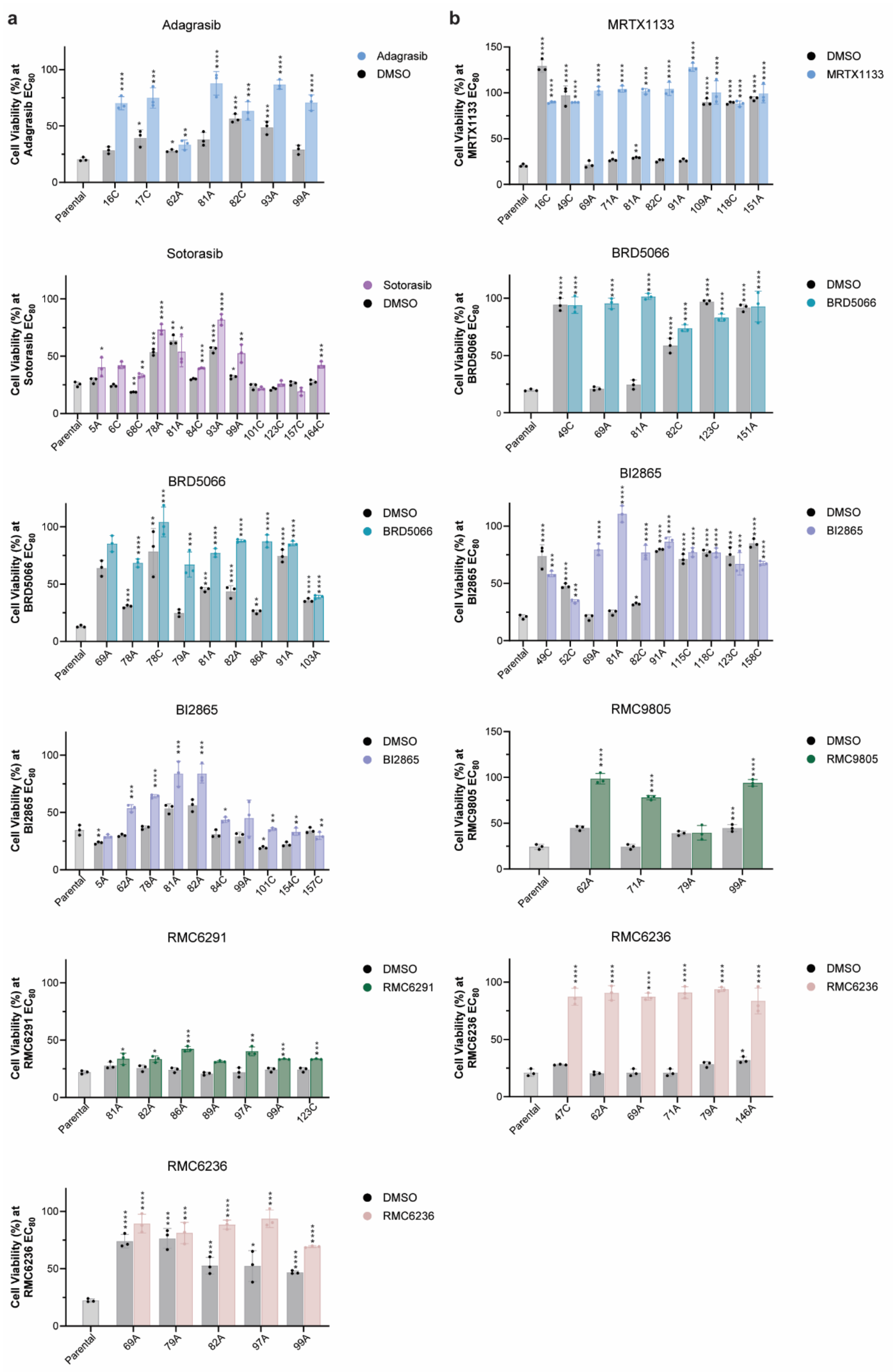


**Figure S9. Delivery of individual sgRNA/BE pairing yields KRAS-resistant cells.** Bar graphs showing inhibitor sensitivity at respective drug selection concentration for $KRAS^{G12C}$ (**a**) and $KRAS^{G12D}$ (**b**) cells transduced with individual sgRNA. Bars and error bars represent the mean and standard deviation, respectively, of measured cell viability (%) of three independent replicates. Statistical significance compared to parental was determined by one-way ANOVA (* p < 0.05, ** p < 0.01, *** p < 0.001, **** p < 0.0001).

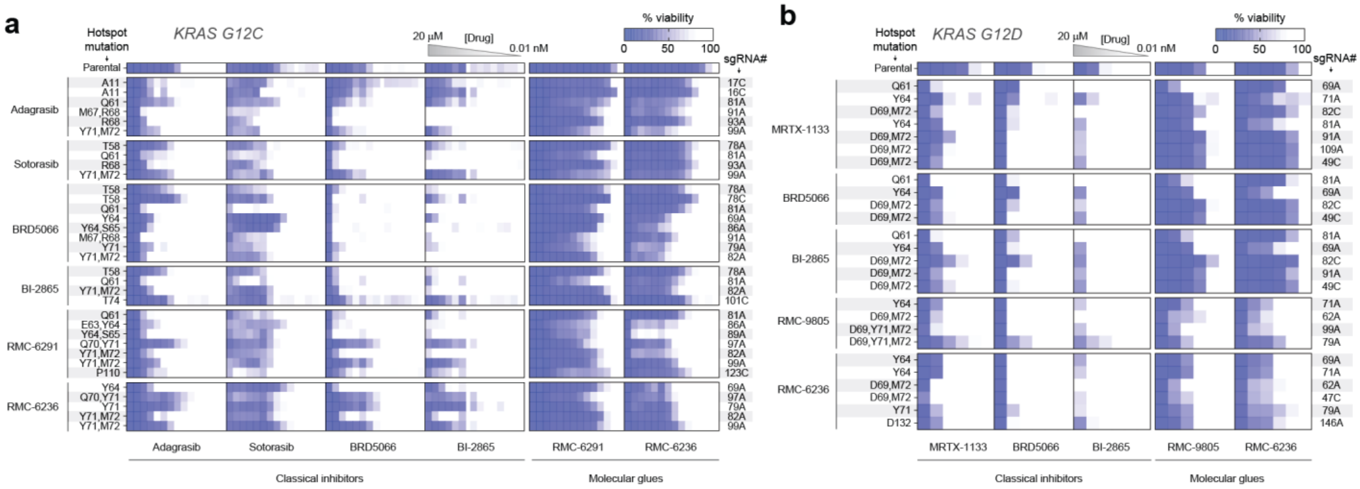


**Figure S10. Viability-matrix with validated sgRNAs confirms drug-resistant KRAS variants. a** Heatmap displaying inhibitor sensitivity of $KRAS^{G12C}$ cells transduced with different sgRNA and base editor pairings. sgRNAs that exhibited a >2 $\log_2$ fold-change versus the library in the screen were introduced into MIA PaCa-2 cells ($KRAS^{G12C}$) along with their respective base editor. Cell lines were tested for sensitivity across all classical inhibitors and molecular glues in a 72-hour cell viability assay using a 12-point, 3-fold serial dilution, with a maximum concentration of 20 μM. Rows represent cells from different drug selections, while columns indicate drugs evaluated in the viability matrix assay. Each heatmap box displays residual cell viability from a single-dose treatment, averaged across three independent replicates. Sites with >10% mutational frequency for a given sgRNA are marked as hotspot mutations. sgRNA# refers to the sgRNA_ID shown in Data S1. Thereby, A and C indicate sgRNAs delivered along with ABE or CBE, respectively. **b** Same as in **a** but for $KRAS^{G12D}$ drug selected cell lines. Drugs were tested in a 72-hour cell viability assay using an 8-point, 3-fold serial dilution, with a maximum concentration of 20 μM.

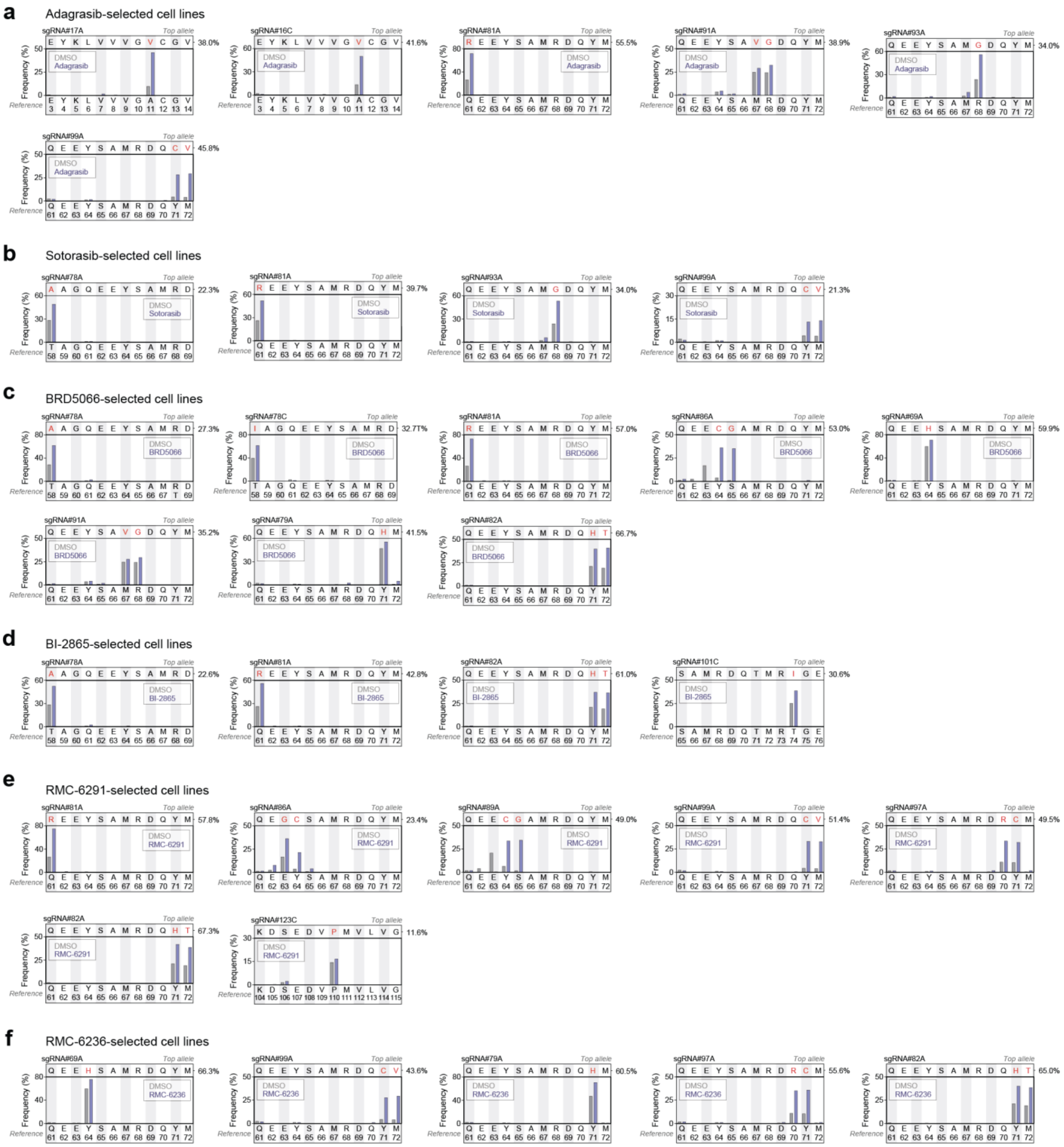

**Figure S11. Genotyping of KRAS$^{G12C}$ drug-selected cells transduced with validated sgRNAs.** After sgRNA transduction, cells were treated either with vehicle (DMSO) or adagrasib (**a**), sotorasib (**b**), BRD5066 (**c**), BI-2865 (**d**), RMC-6291 (**e**), and RMC-6236 (**f**) for 4 weeks to generate drug-resistant cell lines. After drug selection, genomic DNA was extracted, and the targeted exon was PCR-amplified and subjected to deep sequencing. Mutational frequency was calculated as the sum of all alleles carrying a missense mutation at the indicated site. The reference sequence is shown at the bottom, while the most abundant mutated allele (most enriched editing outcome) and its relative frequency are shown at the top. sgRNA# refers to the sgRNA_ID shown in Data S1. Thereby, A and C indicate sgRNAs delivered along with ABE or CBE, respectively.

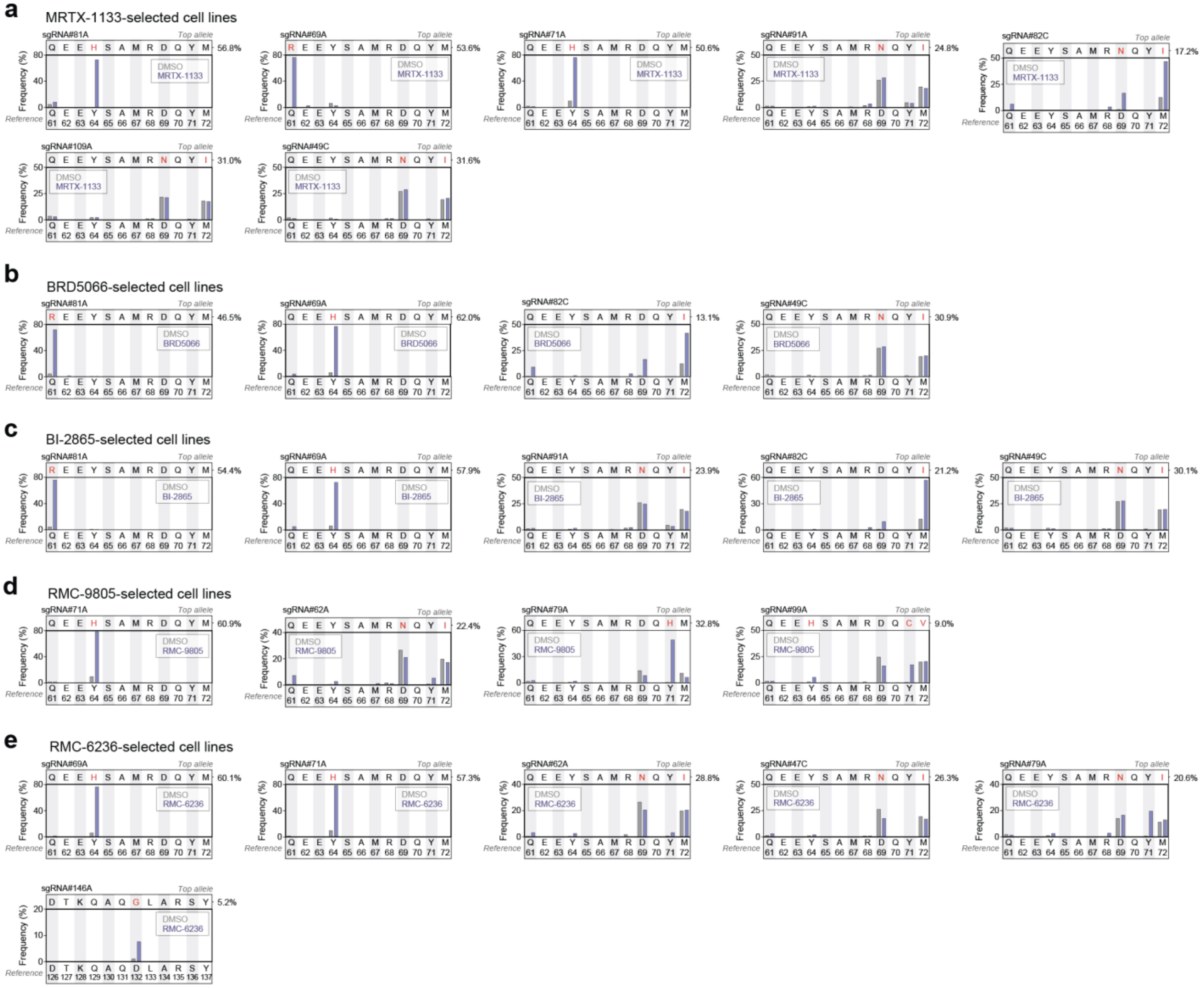


**Figure S12. Genotyping of KRAS$^{G12D}$ drug-selected cells transduced with validated sgRNAs.** Same as Figure S9 but for sgRNAs introduced with their respective base editors into AGS (KRAS$^{G12D}$) cells. Cells were selected with MRTX-1133 (**a**), BRD5066 (**b**), BI-2865 (**c**), RMC-9805 (**d**), and RMC-6236 (**e**).

**Table S1. Sanger sequencing of KRAS4b exons.** *KRAS4b* sequences were analyzed in AGS and MIA PaCa-2 cells to confirm sgRNA library coverage. The top section of the table lists gene-specific primer sequences for each exon, while Sanger sequencing results are shown below. Sequences were trimmed to the *KRAS* coding region, with identified point mutations highlighted in red. Apart from the somatic Gly12 mutation, AGS cells carried only a single silent mutation in exon 4. MIA PaCa2 only had the somatic Gly12 mutation.

| Amplicon description | Direction | Oligonucleotide Sequence (5' → 3') | Source |
|---|---|---|---|
| Exon1 (Round 1) | Forward | ACCCTGACATACTCCCAAGGA | Genewiz |
| | Reverse | GTGCTACAGGGTGTAGAGCA | Genewiz |
| Exon2 (Round 2) | Forward | CCGATGCAGTCTGGAGCAAG | Genewiz |
| | Reverse | AGACTGTGTTTCTCCCTTCTCA | Genewiz |
| Exon3 (Round 2) | Forward | ACCAAAGCCAAAAGCAGTACC | Genewiz |
| | Reverse | AGTTGTGGACAGGTTTTGAAAGA | Genewiz |
| Exon4 (Round 2) | Forward | AACTGCATGCACCAAAAGCC | Genewiz |
| | Reverse | CCTGTACACATGAAGCCATCG | Genewiz |

| Cell line | Amplicon | KRAS coding sequence (5' → 3') |
|---|---|---|
| AGS (KRAS$^{G12D}$) | Exon1 | ATGACTGAATATAAACTTGTGGTAGTTGGAGCTGATGGCGTAGGCAAGAGTGCCTTGACGATACAGCTAATTCAG AATCATTTTGTGGACGAATATGATCCAACAATAGAG |
| | Exon2 | GATTCCTACAGGAAGCAAGTAGTAATTGATGGAGAAACCTGTCTCTTGGATATTCTCGACACAGCAGGTCAAGAG GAGTACAGTGCAATGAGGGACCAGTACATGAGGACTGGGGAGGGCTTTCTTTGTGTATTTGCCATAAATAATACT AAATCATTTGAAGATATTCACCATTATAG |
| | Exon3 | AGAACAAATTAAAAGAGTTAAGGACTCTGAAGATGTACCTATGGTCCTAGTAGGAAATAAATGTGATTTGCCTTCT AGAACAGTAGACACAAAACAGGCTCAGGACTTAGCAAGAAGTTATGGAATTCCTTTTATTGAAACATCAGCAAAGA CAAGACAG |
| | Exon4 | GGTGTTGATGATGCCTTCTATACATTAGTTCGAGAAATTCGAAAACATAAAGAAAAGATGAGCAAAGACGGTAAAA AGAAGAAAAAGAAGTCAAAGACAAAGTGTGTAATTATGTAA |
| MIA PaCa2 (KRAS$^{G12C}$) | Exon1 | ATGACTGAATATAAACTTGTGGTAGTTGGAGCTTGTGGCGTAGGCAAGAGTGCCTTGACGATACAGCTAATTCAG AATCATTTTGTGGACGAATATGATCCAACAATAGAG |
| | Exon2 | GATTCCTACAGGAAGCAAGTAGTAATTGATGGAGAAACCTGTCTCTTGGATATTCTCGACACAGCAGGTCAAGAG GAGTACAGTGCAATGAGGGACCAGTACATGAGGACTGGGGAGGGCTTTCTTTGTGTATTTGCCATAAATAATACT AAATCATTTGAAGATATTCACCATTATAG |
| | Exon3 | AGAACAAATTAAAAGAGTTAAGGACTCTGAAGATGTACCTATGGTCCTAGTAGGAAATAAATGTGATTTGCCTTCT AGAACAGTAGACACAAAACAGGCTCAGGACTTAGCAAGAAGTTATGGAATTCCTTTTATTGAAACATCAGCAAAGA CAAGACAG |
| | Exon4 | GGTGTTGATGATGCCTTCTATACATTAGTTCGAGAAATTCGAAAACATAAAGAAAAGATGAGCAAAGATGGTAAAA AGAAGAAAAAGAAGTCAAAGACAAAGTGTGTAATTATGTAA |

**Table S2. Drug dosing regimen for base editor screening.** A drug dose regimen was established based on parental cell line sensitivity. Cells were subjected to the drug concentration corresponding to an $EC_{30}$ (i.e., 30% growth inhibition) from weeks 0 to 2. Drug concentration was increased to 50% growth inhibition ($EC_{50}$) between weeks 2 and 4 and escalated to the highest concentration corresponding to 80-90% growth inhibition ($EC_{80-90}$) for weeks 4 to 6. If the drug concentration exceeded 5000 nM, the dose for 80% inhibition was used instead to minimize excessive drug exposure and potential off-target effects.

| **Cell line** | **Drug** | **Dose during the indicated treatment period (nM)** | | |
|---|---|---|---|---|
| | | Weeks 0 to 2 | Weeks 2 to 4 | Weeks 4 to 6 |
| MIA PaCa2 ($KRAS^{G12C}$) | Adagrasib | 6.9 | 24 | 183 |
| | Sotorasib | 2.3 | 4.6 | 14.0 |
| | BRD5066 | 71 | 300 | 3200 |
| | BI-2865 | 75 | 163 | 580 |
| | RMC-6291 | 0.1 | 0.16 | 0.4 |
| | RMC-6236 | 2.3 | 4.6 | 14.0 |
| AGS ($KRAS^{G12D}$) | MRTX-1133 | 3.5 | 8.6 | 91.0 |
| | BRD5066 | 35 | 100 | 1510 |
| | BI-2865 | 280 | 610 | 3000 |
| | RMC-9805 | 8.4 | 24 | 370 |
| | RMC-6236 | 1.5 | 4.2 | 66 |

**Table S3. Primers used for deep sequencing of KRAS4b exons.** Amplicons for deep sequencing were generated through two rounds of PCR. In the first round, *KRAS4b* exonic regions were amplified using gene-specific primers with common overhangs (gene-annealing sequences shown in purple). In the second round, Illumina adapter sequences and dual 8-nt indexes (shown in orange) were added for sample demultiplexing. All oligonucleotides were ordered from Sigma or Genewiz.

| Amplicon description | Direction | Oligonucleotide Sequence (5' → 3') | Source |
|---|---|---|---|
| Exon1 (Round 1) | Forward | TCGTCGGCAGCGTCAGATGTGTATAAGAGACAGGGTGGAGTATTTGATAGTGTATTAACC | Sigma |
| | Reverse | GTCTCGTGGGCTCGGAGATGTGTATAAGAGACAGGTATCAAAGAATGGTCCTGCACC | Sigma |
| Exon2 (Round 1) | Forward | TCGTCGGCAGCGTCAGATGTGTATAAGAGACAGCCAGACTGTGTTTCTCCCTTC | Sigma |
| | Reverse | GTCTCGTGGGCTCGGAGATGTGTATAAGAGACAGCTATAATTACTCCTTAATGTCAGC | Sigma |
| Exon3 (Round 1) | Forward | TCGTCGGCAGCGTCAGATGTGTATAAGAGACAGGATATTTGTGTTACTAATGACTGTGC | Sigma |
| | Reverse | GTCTCGTGGGCTCGGAGATGTGTATAAGAGACAGGACATAACAGTTATGATTTTGCAG | Sigma |
| Exon4 (Round 1) | Forward | TCGTCGGCAGCGTCAGATGTGTATAAGAGACAGCATGAAGCCATCGTATATATTCAC | Sigma |
| | Reverse | GTCTCGTGGGCTCGGAGATGTGTATAAGAGACAGCCACTTGTACTAGTATGCCTTAAG | Sigma |
| NGS Indexing (Round 2) | Forward | AATGATACGGCGACCACCGAGATCTACACTAGATCGCTCGTCGGCAGCGTC | Genewiz |
| | Reverse | CAAGCAGAAGACGGCATACGAGATTCGCCTTAGTCTCGTGGGCTCGG | Genewiz |
| | Forward | AATGATACGGCGACCACCGAGATCTACACCTCTCTATTCGTCGGCAGCGTC | Genewiz |
| | Reverse | CAAGCAGAAGACGGCATACGAGATCTAGTACGGTCTCGTGGGCTCGG | Genewiz |
| | Forward | AATGATACGGCGACCACCGAGATCTACACTATCCTCTTCGTCGGCAGCGTC | Genewiz |
| | Reverse | CAAGCAGAAGACGGCATACGAGATTTCTGCCTGTCTCGTGGGCTCGG | Genewiz |
| | Forward | AATGATACGGCGACCACCGAGATCTACACAGAGTAGATCGTCGGCAGCGTC | Genewiz |
| | Reverse | CAAGCAGAAGACGGCATACGAGATGCTCAGGAGTCTCGTGGGCTCGG | Genewiz |
| | Forward | AATGATACGGCGACCACCGAGATCTACACGTAAGGAGTCGTCGGCAGCGTC | Genewiz |
| | Reverse | CAAGCAGAAGACGGCATACGAGATAGGAGTCCGTCTCGTGGGCTCGG | Genewiz |
| | Forward | AATGATACGGCGACCACCGAGATCTACACACTGCATATCGTCGGCAGCGTC | Genewiz |
| | Reverse | CAAGCAGAAGACGGCATACGAGATCATGCCTAGTCTCGTGGGCTCGG | Genewiz |
| | Forward | AATGATACGGCGACCACCGAGATCTACACAAGGAGTATCGTCGGCAGCGTC | Genewiz |
| | Reverse | CAAGCAGAAGACGGCATACGAGATGTAGAGAGGTCTCGTGGGCTCGG | Genewiz |
| | Forward | AATGATACGGCGACCACCGAGATCTACACCTAAGCCTTCGTCGGCAGCGTC | Genewiz |
| | Reverse | CAAGCAGAAGACGGCATACGAGATCCTCTCTGGTCTCGTGGGCTCGG | Genewiz |
| | Forward | AATGATACGGCGACCACCGAGATCTACACGCGTAAGATCGTCGGCAGCGTC | Genewiz |
| | Reverse | CAAGCAGAAGACGGCATACGAGATAGCGTAGCGTCTCGTGGGCTCGG | Genewiz |

**Table S4. Oligonucleotides for validating sgRNAs.** Sequences of sgRNAs that were used for validation are shown. The number corresponds to the sgRNA ID indicated in Data S1. The 20-nt region highlighted in orange in the oligonucleotide sequence corresponds to the protospacer. The flanking sequences were used for Gibson Assembly into the ABE (Addgene 179099) and CBE (Addgene 179096) vectors.

| sgRNA# | Protospacer | Oligonucleotide Sequence (5' → 3') |
|---|---|---|
| 5 | ACTGAATATAAACTTGTGGT | GGAAAGGACGAAACACCGACTGAATATAAACTTGTGGTGTTTGAGAGCTAGAAATAGCAAGTTCAAATAAGGC |
| 6 | GAATATAAACTTGTGGTAGT | GGAAAGGACGAAACACCGGAATATAAACTTGTGGTAGTGTTTGAGAGCTAGAAATAGCAAGTTCAAATAAGGC |
| 16 | TGGAGCTGGTGGCGTAGGCA | GGAAAGGACGAAACACCGTGGAGCTGGTGGCGTAGGCAGTTTGAGAGCTAGAAATAGCAAGTTCAAATAAGGC |
| 17 | GAGCTGGTGGCGTAGGCAAG | GGAAAGGACGAAACACCGGAGCTGGTGGCGTAGGCAAGGTTTGAGAGCTAGAAATAGCAAGTTCAAATAAGGC |
| 21 | TGGCGTAGGCAAGAGTGCCT | GGAAAGGACGAAACACCGTGGCGTAGGCAAGAGTGCCTGTTTGAGAGCTAGAAATAGCAAGTTCAAATAAGGC |
| 47 | ATTACTACTTGCTTCCTGTA | GGAAAGGACGAAACACCGATTACTACTTGCTTCCTGTAGTTTGAGAGCTAGAAATAGCAAGTTCAAATAAGGC |
| 49 | TCCCTTCTCAGGATTCCTAC | GGAAAGGACGAAACACCGTCCCTTCTCAGGATTCCTACGTTTGAGAGCTAGAAATAGCAAGTTCAAATAAGGC |
| 49 | TCCCTTCTCAGGATTCCTAC | GGAAAGGACGAAACACCGTCCCTTCTCAGGATTCCTACGTTTGAGAGCTAGAAATAGCAAGTTCAAATAAGGC |
| 52 | CAGGATTCCTACAGGAAGCA | GGAAAGGACGAAACACCGCAGGATTCCTACAGGAAGCAGTTTGAGAGCTAGAAATAGCAAGTTCAAATAAGGC |
| 56 | GGAAGCAAGTAGTAATTGAT | GGAAAGGACGAAACACCGGGAAGCAAGTAGTAATTGATGTTTGAGAGCTAGAAATAGCAAGTTCAAATAAGGC |
| 62 | TGCTGTGTCGAGAATATCCA | GGAAAGGACGAAACACCGTGCTGTGTCGAGAATATCCAGTTTGAGAGCTAGAAATAGCAAGTTCAAATAAGGC |
| 62 | TGCTGTGTCGAGAATATCCA | GGAAAGGACGAAACACCGTGCTGTGTCGAGAATATCCAGTTTGAGAGCTAGAAATAGCAAGTTCAAATAAGGC |
| 68 | GTACTCCTCTTGACCTGCTG | GGAAAGGACGAAACACCGGTACTCCTCTTGACCTGCTGGTTTGAGAGCTAGAAATAGCAAGTTCAAATAAGGC |
| 69 | CTGTACTCCTCTTGACCTGC | GGAAAGGACGAAACACCGCTGTACTCCTCTTGACCTGCGTTTGAGAGCTAGAAATAGCAAGTTCAAATAAGGC |
| 71 | GCACTGTACTCCTCTTGACC | GGAAAGGACGAAACACCGGCACTGTACTCCTCTTGACCGTTTGAGAGCTAGAAATAGCAAGTTCAAATAAGGC |
| 78 | CGACACAGCAGGTCAAGAGG | GGAAAGGACGAAACACCGCGACACAGCAGGTCAAGAGGGTTTGAGAGCTAGAAATAGCAAGTTCAAATAAGGC |
| 78 | CGACACAGCAGGTCAAGAGG | GGAAAGGACGAAACACCGCGACACAGCAGGTCAAGAGGGTTTGAGAGCTAGAAATAGCAAGTTCAAATAAGGC |
| 79 | GTACTGGTCCCTCATTGCAC | GGAAAGGACGAAACACCGGTACTGGTCCCTCATTGCACGTTTGAGAGCTAGAAATAGCAAGTTCAAATAAGGC |
| 81 | GCAGGTCAAGAGGAGTACAG | GGAAAGGACGAAACACCGGCAGGTCAAGAGGAGTACAGGTTTGAGAGCTAGAAATAGCAAGTTCAAATAAGGC |
| 82 | CTCATGTACTGGTCCCTCAT | GGAAAGGACGAAACACCGCTCATGTACTGGTCCCTCATGTTTGAGAGCTAGAAATAGCAAGTTCAAATAAGGC |
| 82 | CTCATGTACTGGTCCCTCAT | GGAAAGGACGAAACACCGCTCATGTACTGGTCCCTCATGTTTGAGAGCTAGAAATAGCAAGTTCAAATAAGGC |
| 84 | AAGAGGAGTACAGTGCAATG | GGAAAGGACGAAACACCGAAGAGGAGTACAGTGCAATGGTTTGAGAGCTAGAAATAGCAAGTTCAAATAAGGC |
| 86 | GAGGAGTACAGTGCAATGAG | GGAAAGGACGAAACACCGGAGGAGTACAGTGCAATGAGGTTTGAGAGCTAGAAATAGCAAGTTCAAATAAGGC |
| 89 | GTACAGTGCAATGAGGGACC | GGAAAGGACGAAACACCGGTACAGTGCAATGAGGGACCGTTTGAGAGCTAGAAATAGCAAGTTCAAATAAGGC |
| 91 | TGCAATGAGGGACCAGTACA | GGAAAGGACGAAACACCGTGCAATGAGGGACCAGTACAGTTTGAGAGCTAGAAATAGCAAGTTCAAATAAGGC |
| 93 | AATGAGGGACCAGTACATGA | GGAAAGGACGAAACACCGAATGAGGGACCAGTACATGAGTTTGAGAGCTAGAAATAGCAAGTTCAAATAAGGC |
| 97 | GACCAGTACATGAGGACTGG | GGAAAGGACGAAACACCGGACCAGTACATGAGGACTGGGTTTGAGAGCTAGAAATAGCAAGTTCAAATAAGGC |
| 99 | CAGTACATGAGGACTGGGGA | GGAAAGGACGAAACACCGCAGTACATGAGGACTGGGGAGTTTGAGAGCTAGAAATAGCAAGTTCAAATAAGGC |
| 99 | CAGTACATGAGGACTGGGGA | GGAAAGGACGAAACACCGCAGTACATGAGGACTGGGGAGTTTGAGAGCTAGAAATAGCAAGTTCAAATAAGGC |
| 101 | GGACTGGGGAGGGCTTTCTT | GGAAAGGACGAAACACCGGGACTGGGGAGGGCTTTCTTGTTTGAGAGCTAGAAATAGCAAGTTCAAATAAGGC |
| 102 | ACTGGGGAGGGCTTTCTTTG | GGAAAGGACGAAACACCGACTGGGGAGGGCTTTCTTTGGTTTGAGAGCTAGAAATAGCAAGTTCAAATAAGGC |
| 103 | GAGGGCTTTCTTTGTGTATT | GGAAAGGACGAAACACCGGAGGGCTTTCTTTGTGTATTGTTTGAGAGCTAGAAATAGCAAGTTCAAATAAGGC |

| | | |
|---|---|---|
| 109 | AAGATATTCACCATTATAGG | GGAAAGGACGAAACACCGAAGATATTCACCATTATAGGGTTTGAGAGCTAGAAATAGCAAGTTCAAATAAGGC |
| 115 | ATTAAAAGAGTTAAGGACTC | GGAAAGGACGAAACACCGATTAAAAGAGTTAAGGACTCGTTTGAGAGCTAGAAATAGCAAGTTCAAATAAGGC |
| 118 | TATTTCCTACTAGGACCATA | GGAAAGGACGAAACACCGTATTTCCTACTAGGACCATAGTTTGAGAGCTAGAAATAGCAAGTTCAAATAAGGC |
| 121 | GACTCTGAAGATGTACCTAT | GGAAAGGACGAAACACCGGACTCTGAAGATGTACCTATGTTTGAGAGCTAGAAATAGCAAGTTCAAATAAGGC |
| 123 | GATGTACCTATGGTCCTAGT | GGAAAGGACGAAACACCGGATGTACCTATGGTCCTAGTGTTTGAGAGCTAGAAATAGCAAGTTCAAATAAGGC |
| 133 | TAAGTCCTGAGCCTGTTTTG | GGAAAGGACGAAACACCGTAAGTCCTGAGCCTGTTTTGGTTTGAGAGCTAGAAATAGCAAGTTCAAATAAGGC |
| 135 | TTCTTGCTAAGTCCTGAGCC | GGAAAGGACGAAACACCGTTCTTGCTAAGTCCTGAGCCGTTTGAGAGCTAGAAATAGCAAGTTCAAATAAGGC |
| 139 | GTAGACACAAAACAGGCTCA | GGAAAGGACGAAACACCGGTAGACACAAAACAGGCTCAGTTTGAGAGCTAGAAATAGCAAGTTCAAATAAGGC |
| 146 | AGGACTTAGCAAGAAGTTAT | GGAAAGGACGAAACACCGAGGACTTAGCAAGAAGTTATGTTTGAGAGCTAGAAATAGCAAGTTCAAATAAGGC |
| 151 | TTACTTACCTGTCTTGTCTT | GGAAAGGACGAAACACCGTTACTTACCTGTCTTGTCTTGTTTGAGAGCTAGAAATAGCAAGTTCAAATAAGGC |
| 153 | TTATTTCAGTGTTACTTACC | GGAAAGGACGAAACACCGTTATTTCAGTGTTACTTACCGTTTGAGAGCTAGAAATAGCAAGTTCAAATAAGGC |
| 154 | AACATCAGCAAAGACAAGAC | GGAAAGGACGAAACACCGAACATCAGCAAAGACAAGACGTTTGAGAGCTAGAAATAGCAAGTTCAAATAAGGC |
| 157 | TTTTATGTATTTCAGGGTGT | GGAAAGGACGAAACACCGTTTTATGTATTTCAGGGTGTGTTTGAGAGCTAGAAATAGCAAGTTCAAATAAGGC |
| 158 | TATGTATTTCAGGGTGTTGA | GGAAAGGACGAAACACCGTATGTATTTCAGGGTGTTGAGTTTGAGAGCTAGAAATAGCAAGTTCAAATAAGGC |
| 163 | TTTCGAATTTCTCGAACTAA | GGAAAGGACGAAACACCGTTTCGAATTTCTCGAACTAAGTTTGAGAGCTAGAAATAGCAAGTTCAAATAAGGC |
| 164 | GATGATGCCTTCTATACATT | GGAAAGGACGAAACACCGGATGATGCCTTCTATACATTGTTTGAGAGCTAGAAATAGCAAGTTCAAATAAGGC |
| 166 | GCCTTCTATACATTAGTTCG | GGAAAGGACGAAACACCGGCCTTCTATACATTAGTTCGGTTTGAGAGCTAGAAATAGCAAGTTCAAATAAGGC |

**Data S1 Notes. KRAS sgRNA library sequences.** List of protospacer sequences and corresponding oligonucleotides ordered from Twist Bioscience for base editor mutagenesis library construction. Protospacers were designed based on the KRAS4b isoform using the relaxed "NGN" PAM site. The library was constructed for the KRAS4b gene without primary mutations and used for mutagenesis in MIA PaCa-2 ($KRAS^{G12C}$) and AGS ($KRAS^{G12D}$) cells. Intronic sequences were excluded from the sgRNA library design, except for the 20 bp flanking each exon. The ordered oligonucleotides include adapter sequences for cloning and Gibson Assembly into designated vectors and subpool barcode sequences for targeted PCR amplification of specific library fractions. Missense mutations for each base editor were predicted based on sgRNA mapping, assuming that any cytosine (for CBE) or adenine (for ABE) within the +4 to +8 editing window of the protospacer sequence could be converted to thymine or guanine, respectively.

**Data S2 Notes. Base editor screening data in MIA PaCa2 ($KRAS^{G12C}$) cells.** sgRNA counts were expressed in ppm and $log_2$-transformed with a pseudo count of one. The transformed data were normalized to the plasmid library to generate resistance scores, where higher scores indicate resistance, and lower scores reflect drug sensitivity. Enrichment data were analyzed at two, four, and six weeks of selection, with drug doses increased every two weeks. The dosing regimen for each drug is detailed in Table S2. All selections, including vehicle-treated controls, were performed in three independent replicates.

**Data S3 Notes. Base editor screening data in AGS ($KRAS^{G12D}$) cells.** Same as in Data S2 Notes but for AGS cells and $KRAS^{G12D}$.

**Data S4 Notes. Deep sequencing of editing outcomes in base editor screening.** Genomic DNA was extracted after completing 6 weeks of drug selection, and the exons in *KRAS4b* in each selection were analyzed by deep sequencing. Each allele in the table corresponds to a full exon sequenced as a single amplicon, and allele frequencies were determined using CRISPRessoV2 and a custom script. The drug-selection column indicates cell line, base editor, and drug used for the enrichment of drug-resistant cells. The allele column lists detected alleles along with their corresponding translated products. Outcomes from the base editor screen with an allele frequency above 1% are shown. Notably, multiple alleles may produce identical translated proteins due to silent mutations. For representation in figure analyses, identical translated products were consolidated, and their allele frequencies were combined.

**Data S5 Notes. Deep sequencing of editing outcomes for validated sgRNAs.** Same as in Data S4 Notes but for the validated sgRNAs. sgRNAs were delivered to MIA PaCa2 ($KRAS^{G12C}$) or AGS ($KRAS^{G12D}$) cells along with the indicated editor and subjected to drug selection for four weeks at the highest concentration depicted in Table S2. Genomic DNA was extracted, and the exon targeted by the sgRNA was subjected to deep sequencing. Alleles with more than 1% frequency are shown for each sgRNA. Only sgRNAs that conferred drug resistance are shown. The drug selection and deep sequencing were performed once ($n$=1).
